# Role of protein dynamics in enthalpy-driven recognition of topologically distinct dsRNAs by dsRBDs

**DOI:** 10.1101/862326

**Authors:** H Paithankar, J Chugh

## Abstract

Taking leads from the fact that a handful of double-stranded RNA-binding domains (dsRBDs) interact with a massive number of topologically distinct double-stranded RNAs (dsRNAs) in crucial biological pathways, and to understand the adaptability required by dsRBDs to target the pool of dsRNA substrates, we employed two independent model dsRBDs in an ITC and NMR spectroscopy based study. Our previous study revealed the presence of microsecond timescale dynamics in RNA-binding regions in the two dsRBDs studied. In the current study, results from ITC-based titrations showed that the binding of dsRBD with topologically distinct dsRNAs is enthalpy-driven, with each dsRNA-dsRBD pair having distinct combination of enthalpy-entropy yielding a similar change in free energy upon RNA-binding. We also show that the each of the dsRNA, used in this study, binds to the dsRBD in a unique mode. Further, intrinsic conformational exchange present in the RNA-binding regions of the apo-dsRBD was shown to quench upon binding with a dsRNA, while conformational exchange got induced at the residues that are in close proximity to where exchange was present in the apo-protein. This apparent relay of conformational exchange from one site to the other site upon dsRNA-binding thus suggests the importance of intrinsic dynamics to adapt to target a variety of dsRNA-shapes.

**Statement of Significance:** This study reports that the interaction between dsRBDs and dsRNAs is enthalpy-driven, is perturbed by subtle changes in dsRNA shapes, and exhibits a classic case of enthalpy-entropy compensation. Further, the intrinsic microsecond timescale conformational exchange present in the apo-dsRBD was observed to get quenched upon RNA-binding. An apparent relay of conformational exchange was also observed from quenched sites to spatially proximal sites upon RNA-binding, suggesting highly adaptive nature of the dsRBD, a critical feature required to target topologically distinct dsRNA.

## Introduction

In addition to the protein-protein and protein-DNA interactions, the interaction between doublestranded RNAs (dsRNAs) and dsRNA-binding domains (dsRBDs) of proteins plays a vital role in complex cellular pathways like small RNA biogenesis (1, 2), post-transcriptional RNA modification (3, 4), RNA transport (5), etc. To perform a diverse array of functions, the dsRBDs need to interact with a wide range of RNA molecules available in the cellular matrix and specifically recognize the target RNA to be processed. The dsRBDs have been reported to target dsRNAs in a non-sequence specific manner and interact with an assortment of dsRNA conformations – ranging from simple A-form helices to as complex as L-shaped tRNA molecules (6–12). Broadly, dsRBDs are 65-70 aa long polypeptide chains with a well-defined α_1_-β_1_-β_2_-β_3_-α_2_ structural fold, where the two α-helices are packed on to side of a plane of three anti-parallel β-strands (7, 13). The middle of helix α_1_ the loop between β_1_ and β_2_ strands, and the N-terminal of helix α_2_ from dsRBDs together constitutes the dsRNA-recognition surface that engages a stretch of ~12 bp long minor-major-minor groves on dsRNA, where the two minor groves interact via ribose sugars and the major grove interacts via phosphate backbone (7, 13). Conformational assortment in target dsRNAs stems due to the frequent occurrence of non-canonical base-pairs leading to the presence of secondary structure breaks in the form of bulges and/or internal loops. The bulges or internal loops are not only catalytically important as they provide active centers (14, 15); they also change relative orientation (defined by the Euler angles (16)) of adjoining A-form helices, thereby changing the spread of minor-major-minor groves along the length of the dsRNA. Therefore, each of such dsRNA conformation presents a unique spread of minor-major-minor groves that is targeted by a particular dsRBD. To effectively and efficiently process these dsRNAs, dsRBDs must recognize the target dsRNA substrate specifically from the cellular milieu.

There are only 22 dsRBD-dsRNA structures available (as of September 2019 in NDB (http://ndbserver.rutgers.edu/) and PDB (https://www.rcsb.org/) database together) in the literature that have been solved with A-form helical RNAs with or without perturbations in the secondary structure (17–20). While, structural studies provide crucial insights into the mode of recognition and binding regions of dsRNA-dsRBD interaction (10–12, 21–24); the dynamics measurements address the vibrant feature of the interaction present in the dsRBDs that possibly allows them to adapt and interact with multiple partners in a shape-dependent manner. Furthermore, tandem dsRBDs have been shown to diffuse along the length of the dsRNA in an ATP-independent manner in search for ‘proper’ substrates (25), thereby suggesting that the dynamic interface is central to dsRNA recognition by dsRBDs. Such dynamics measurements have been reported in proteins (TRBP, ADAR1, ADAR2, and Staufen) with tandem dsRBDs using FRET-based studies and have shown differential degree of sliding motions on dsRNAs of varying length and secondary structure (26, 27).

In the present study, we have attempted to understand the structural basis of dsRNA recognition by dsRBDs by studying interactions between two model dsRBDs and topologically different dsRNAs. Two distinct dsRBDs, namely, dsRBD1 from human TAR RNA binding protein (TRBP2) and *Drosophila* Adenosine deaminase that act on RNA (dADAR) – known to adopt canonical dsRBD fold and target multiple dsRNAs – were employed (6, 25–27). While TRBP was first identified as a protein involved in interaction with the HIV TAR RNA element (28), recent studies have implicated its role in the microRNA (miR) biogenesis where it interacts with precursors of miRs and presents them to Dicer (an RNase III endonuclease) for further processing (29–32). The precursors of miRs (~1917 human precursor-miRs sequences are known till date (33), of which a large number are processed by TRBP) are rarely pure A-form helices and often contain internal loops and bulges thereby creating a large pool of conformational targets for TRBP. A recent report has shown that TRBP exhibits dual recognition mechanism where short binding and long binding events allow to discriminate between substrate dsRNAs from other RNAs in the cellular environment (34). Likewise, ADAR proteins are involved in regulating gene expression by editing the RNA base Adenine to inosine (35–37). While there is an evidence that suggests dsRBD2 of human ADAR2 targets its RNA in a sequence-specific manner by recognizing the base pair 3′ adjacent to the editing site (4); yet another report claims that the interaction between dsRBD1 of *Drosophila* ADAR and dsRNAs is relatively less sequence-specific (6), thereby creating a slight haze on the targeting mechanism of dsRBDs of ADARs.

A recent study from our lab has reported the presence of microsecond timescale dynamics in the two dsRBDs. Based on these results, we have proposed that the conformational ensemble of the dsRBDs might be responsible for the recognition of conformationally distinct target dsRNAs (38). Here, we investigated the binding modes of these dsRBDs with topologically distinct dsRNAs by performing a combination of ITC-based titration and NMR spectroscopy studies. The ITC and NMR data suggest that the binding modes of dsRBD towards topologically distinct dsRNAs are unique. NMR data showed that the motional modes of dsRBDs are perturbed by dsRNAs in a shape-dependent fashion in order to allow efficient binding of dsRBD. Our studies on small RNA construct further showed that the conformational dynamics intrinsic to the dsRBDs fold allow them to bind to the dsRNA substrate via a conformational rearrangement mechanism. These results will pave the way to understand the adaptation required in dsRBDs to target a variety of dsRNA targets while ignoring the helical defects present in such dsRNAs and process them for downstream cellular pathways.

## Materials and Methods

### Protein expression and purification

The TRBP2-dsRBD1 (1-105 aa) and dADAR-dsRBD1 (48-140 aa) proteins were overexpressed and purified as described elsewhere (6, 39). For ^15^N labeling, the proteins were prepared by growing the *E. coli* BL21(*DE3*) cells in the M*9* media containing the ^15^NH_4_Cl (Cambridge Isotope Laboratories, USA) as a sole source of nitrogen.

### RNA sequence design and sample preparation

The RNA sequences were designed based on the fact that the TRBP2-dsRBD1 is known to interact with miR-16-1 duplex (9). The miR-16-1 duplex contains a bulge (created by an unpaired uridine), and an internal loop (created by a A•A mismatch) in the dsRNA-binding region (40), thereby creating a deviation from the A-form helical structure (Figure 1A). The passenger strand of miR-16-1 was mutated to generate the following three RNA sequence mutants: i) miR-16-1-M with only the mismatch (Figure 1B); ii) miR-16-1-B with only the bulge (Figure 1C); and iii) miR-16-1-D which forms a perfect duplex by removing both the bulge and the internal loop (Figure 1D). A shorter duplex RNA oligo (D10RNA) was also designed from miR-16-1-D, keeping a minimum length of RNA (10 bp), that can potentially interact with dsRBD (41). All the sequences have been listed in the Supplementary Table S1.

**Figure 1:**
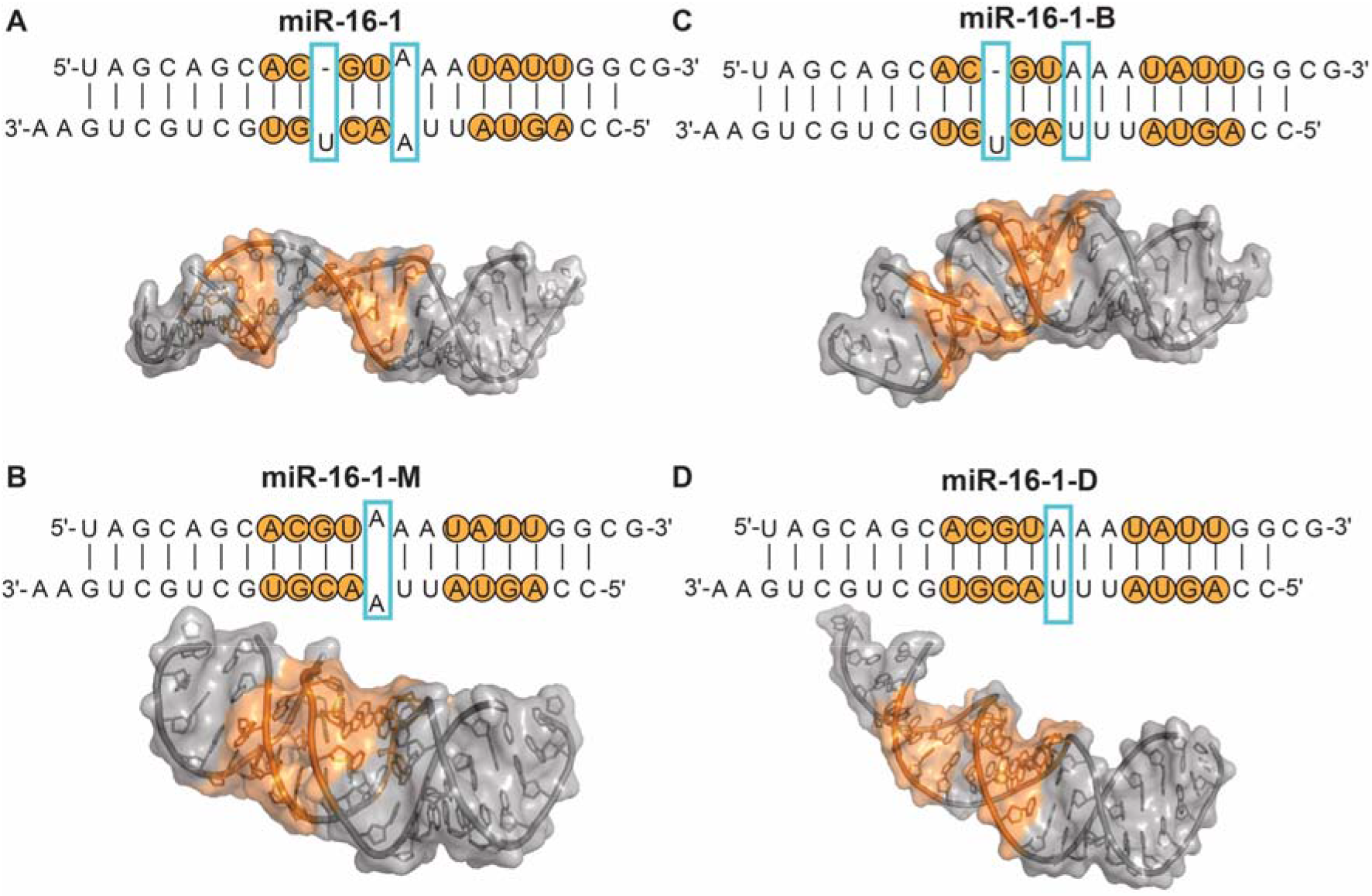
dsRNA sequences designed to test shape-dependent binding with dsRBDs. Four dsRNA sequences designed based on (A) miR-16-1 sequence containing a bulged Uridine and a A-A mismatch, a known binder of TRBP2-dsRBD1; (B) bulged Uridine is deleted and only mismatch is preserved; (C) A-A mismatch is concerted to a stable A-U base pair, while the bulged Uridine is preserved; and (D) bulged Uridine is deleted, and A-A mismatch is converted into a stable A-U base pair. dsRBD-binding site has been marked in orange and helical imperfection has been marked in cyan boxes. Simulations showing topological perturbations due to deletion/mutation has been indicated using the tertiary structure obtained using SimRNA (42, 43).

All RNAs were procured from either Integrated DNA Technologies (Coralville, Iowa, USA) or Sigma Aldrich (St. Louis, Missouri, USA) as single strands. RNA annealing was carried out by mixing the respective strands in 1:1 ratio. The strands were denatured by heating at 95°C for 5 min and then allowed to anneal at 4°C for 10 min. Annealing of the two strands was confirmed by ‘H NMR by the presence of imino proton peaks (data not shown). The annealed samples were maintained in the buffer A (10 mM Sodium phosphate, 100 mM NaCl, 1 mM EDTA, 1 mM DTT), pH 6.4 for performing ITC and NMR-based studies.

### RNA 3D structure modeling

The 3D structure for the four RNA sequences, based on miR-16-1, was modeled using SimRNA (42, 43) that uses a coarse-grained simulation, relies on Monte Carlo method for sampling the conformational space, and further employs a statistical potential to approximate the energy to identify the low-energy conformations. The secondary structure of the RNA sequences for the input to the SimRNA program was obtained from DINAMelt (44, 45).

### Isothermal Titration Calorimetry

All isothermal titration calorimetry experiments were performed on MicroCal iTC-200 at 25°C. The protein and RNA samples were diluted to the required concentrations using buffer A. Briefly, 30 μM protein (TRBP2-dsRBD1 or ADAR-dsRBD1) was added in the cell while RNA (as ligand) was present in the syringe. For different RNA constructs, the molar ratio of protein to RNA was optimized to obtain a saturated calorimetric signal at the end of the binding studies. The heat of dilution was determined by titrating the RNA into the buffer, buffer into the protein, and buffer into the buffer. In each injection, 2 μl of RNA was added to the sample cell over 4 seconds with interval of 120 seconds between the injections. The data obtained was corrected for the heat of dilution and then fitted to the one set of sites model in Origin 7.0 provided with the instrument. The RNA concentration used in the syringe for the four RNAs are miR-16-1 = 30 μM; miR-16-1-D = 120 μM; miR-16-1-M = 300 μM; miR-16-1-B = 150 μM; and D10RNA = 240 μM; respectively.

### NMR spectroscopy

All the NMR experiments were carried out on 1) Bruker 600 MHz NMR spectrometer equipped with quadruple-resonance (^1^H/^15^N/^13^C/^31^P) 5 mm Cryoprobe, X, Y, Z-gradients, and dual receiver operating (located in-house); 2) Bruker 600 MHz NMR spectrometer equipped with TCI ^13^C-enhanced 5 mm cryoprobe, Z-gradient, and deuterium decoupling (located at NCBS Bangalore, India); and 3) Bruker 950 MHz NMR spectrometer equipped with TCI cryoprobe, Z-gradient, and deuterium decoupling (located at Institute of Protein Research, Osaka University, Osaka, Japan). Unless mentioned otherwise, all the NMR experiments have been recorded at 298 K. All the NMR spectra have been processed using NMRPipe/NMRDraw (46) and analyzed using SPARKY (47).

^1^H-^15^N-HSQC was measured on TRBP2-dsRBD1 in the absence of the RNA, and in increasing concentrations of RNAs from 0.05 to 0.25 equivalents (0.05 to 2.5 equivalents in case of D10RNA) of the protein. The change in the intensities were plotted as a function of concentration of the RNA added to the protein. The chemical shift perturbations (Δδ) were calculated as (48):

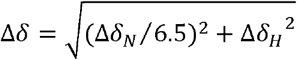

where, Δδ_N_ and Λδ_H_ represent the change in the ^15^N and ^1^H chemical shifts in the ^1^H-^15^N-HSQC spectrum of ^15^N-TRBP2-dsRBD1 upon addition of 0.1 equivalents of RNA (0.5 equivalents of D10RNA), respectively.

Nuclear spin relaxation experiments were measured on 600 MHz NMR spectrometer on a 200 μM ^15^N-TRBP2-dsRBD1 sample in complex with RNAs at 1:0.5 protein:RNA ratio in Shigemi tubes. For D10RNA-bound TRBP2-dsRBD1, ^15^N transverse relaxation rates (*R*_2_) were measured with 8 CPMG delays of 17, 34*, 51, 68, 85, 102, 136*, and 170 ms with a CPMG loop length of 17 ms. ^15^N longitudinal relaxation rates (*R*_1_) were measured with 8 inversion recovery delays of 10, 30, 50*, 100, 200, 350, 500*, and 700 ms for RNA-bound state. Delays marked with an asterisk in both the experiments were measured in duplicate for estimation of errors in relaxation rates. Steady-state ^1^H-^15^N heteronuclear nOe measurements were carried out with a ^1^H saturation time of 3 s and a relaxation delay of 2 s for both the proteins. For the experiment without ^1^H saturation, the relaxation delay of 5 s was used. All the nuclear spin relaxation experiments were measured in an interleaved fashion and with randomized order of delays.

^15^N relaxation dispersion experiments were measured using constant time CPMG (Carr-Purcell-Meiboom-Gill) experiments (49) on 600 MHz NMR spectrometer. For D10RNA-bound TRBP2-dsRBD1, each relaxation dispersion profile was composed of 9 points with ν_cpmg_ values of 25, 50*, 100, 200, 350, 500, 650*, 800, and 1000 Hz at constant relaxation time, T_relax_, of 40 ms. Duplicate points were measured for error analysis and have been marked with an asterisk.

Heteronuclear Adiabatic Relaxation Dispersion (HARD) experiments (50, 51) on D10RNA-bound TRBP2-dsRBD1 were performed at 600 MHz NMR spectrometer. Composite adiabatic pulse of 16 ms containing four hyperbolic secant family of pulses of 4 ms each were used for spin-locking the ^15^N magnetization in both R_1ρ_ and R_2ρ_ experiments, as described earlier (50, 51). Relaxation dispersion was created with adiabatic hyperbolic secant pulses with different stretching factors (n=1,2,4,6,8). As the stretching factor was increased from 1 to 8, the effective spin-lock field strength increased. Relaxation delays for adiabatic R_1ρ_ and R_2ρ_ experiments were varied by varying the number of composite pulses applied during evolution. The relaxation delays used were 0, 16, 32, 48 and 64 ms corresponding to the number of composite adiabatic pulses of 0, 1, 2, 3, 4. R_1_ experiments were acquired in the same way as R_1ρ_ and R_2ρ_ experiments without using the adiabatic pulse during evolution. Relaxation delays used for the R_1_ experiment were 16, 48, 96, 192, 320, 480 and 640 ms. An inter-scan delay of 2.5 s was used in all the above relaxation experiments.

### NMR relaxation data analysis

NMR relaxation data analysis for the D10RNA-bound TRBP2-dsRBD1 was carried out in a similar way as was done for apo-TRBP2-dsRBD1 described elsewhere (38).

Relaxation rates in *R*_1_/*R*_2_/*R*_1ρ_/*R*_2ρ_ experiments were calculated by fitting the intensity data against relaxation delays to mono-exponential decays in Mathematica (52). Errors in the relaxation rates were calculated using a combination of duplicate delay data and Monte Carlo simulations. [^1^H]-^15^N nOe values were obtained as a ratio of the intensity of respective peaks of the spectra recorded with and without saturation. Errors in nOe values were obtained by propagating the errors from RMSD values of baseline noise as obtained from Sparky.

In CPMG relaxation dispersion experiments, effective transverse relaxation rates at each CPMG frequency were extracted using the following equation (49, 53):

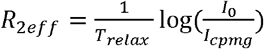

where, I_0_ is the intensity of a peak in reference spectrum and I_cpmg_ is the intensity of the respective peak in the spectrum with CPMG field applied at a frequency ν_cpmg_. The R_2eff_ rates thus obtained were plotted against the CPMG frequency ν_cpmg_.

In HARD experiments, the relaxation rates (*R*_1_, *R*_1ρ_, and *R*_2ρ_) along with rotational correlation time calculated from quadric_diffusion program (54) was used to fit the data to the time-dependent Bloch-McConnell equation assuming a two-site exchange model as was done for the apo-state of the protein (38, 51). The solution to the equation by using geometric approximation method allowed to extract the dynamic parameters (the conformational exchange rate, *k*_ex_, the equilibrium populations of the two states, p_A_ or p_B_, and the chemical shift difference, Δω) and the relaxation rates during the chemical exchange process as described in detail elsewhere (51).

## Results

### Thermodynamics of dsRBD-dsRNA interactions

Four dsRNAs (Figure 1), designed based on miR-16-1 – a known binder of TRBP2-dsRBD1 (9), were subjected to simulation using SimRNA (42, 43) to highlight the topological differences in the conformations arising due to the presence of bulge and/or internal loop (Figure 1). It is important to note from the simulation results that the subtle changes (single-base mutation/deletion) in the RNA-sequence — upon removing either bulged Uridine or removing internal loop (created by the A•A mismatch) by mutating A-to-U — created differences in the overall shape of the RNA. This observation is corroborated with a previous finding where Al-Hashimi and co-workers observed that the global conformation of an RNA is linked to its secondary structure (16).

To explore the effect of differences observed in the global conformation of RNA on the thermodynamics of binding with the partner protein(s), ITC-based binding experiments were performed, where TRBP2-dsRBD1 and dADAR-dsRBD1 were subjected to titrations against all four dsRNAs (Figure 2 and Figure S1). The dissociation constant, K_d_, for TRBP2-dsRBD1 ranged between 0.50±0.04 to 2.12±0.22 μM; while that of dADAR-dsRBD1 ranged between 0.27±0.05 to 0.78±0.24 μM (Table S1). The corresponding changes in ΔH and TΔS ranged from −9.4±0.2 to −34.8±0.6 kcal/mol and from −1.6±0.2 to −26.2±0.6 kcal/mol for TRBP2-dsRBD1; and from −24.6±1.1 to - 80.9±3.2 kcal/mol and from −16.3±0.9 to −72.1±3.1 kcal/mol for dADAR-dsRBD1, respectively. It is intriguing to see that while ΔG of binding for both the proteins and for all four dsRNAs is ~8 kcal/mol, ΔH and TΔS values are distinct for each dsRBD-dsRNA interacting pair. Respective changes in ΔH and TΔS appear like a classic example of enthalpy-entropy compensation (55–57), where any change in ΔH is restored by the corresponding change in TΔS (or *vice versa*), keeping ΔG similar across the changes in dsRNA conformation. Although there are numerous physical factors, including solvent reorganization, structure of the protein and the ligand, their intrinsic flexibility, etc. that contribute to ΔH and TΔS of the binding (55, 58, 59), the distinctness in the thermodynamic parameters obtained from different titrations carried out here may be explained by 1) the changes in global dsRNA conformation from one dsRNA to another (16, 60–62), and 2) conformational entropy present in the two dsRBDs by virtue of ms-μs timescale motions as reported previously by our group (38).

**Figure 2:**
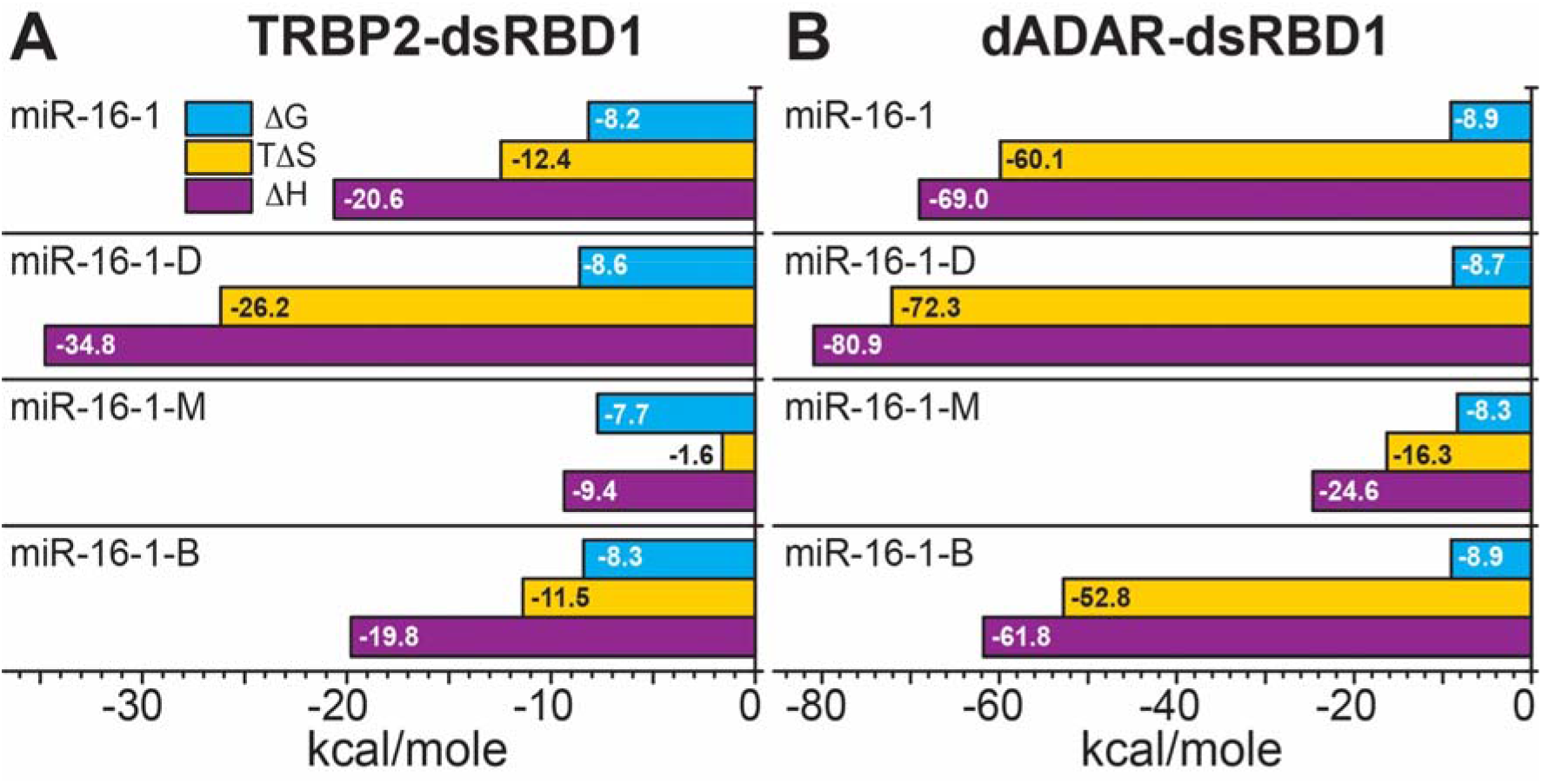
Thermodynamic parameters (ΔG, ΔH, and TΔS) plotted as horizontal bars in the units of kcal/mole as obtained from ITC-based titration of (A) TRBP2-dsRBD1, and of (B) dADAR-dsRBD1 against the four dsRNAs described in figure 1. Color scheme for horizontal bars has been indicated in the figure as ΔG (cyan), TΔS (yellow), and ΔH (purple). A negative value of ΔH is favourable, while that of TΔS is unfavourable for the binding reaction.

Interestingly, the number of binding sites of dsRBDs in the dsRNAs were also found to be different for all four dsRNAs (Table S2). While, the weakest binder dsRNA, miR-16-1-M, to TRBP2-dsRBD1 and dADAR-dsRBD1 had number of protein binding sites as 1.5 and 1.6, respectively, the strongest binding dsRNA, miR-16-1-D for TRBP2-dsRBD1 and miR-16-1-B for dADAR-dsRBD1, had number of binding sites as 5.2 and 4.4, respectively.

### Intermediate-exchange observed in dsRBD-dsRNA interactions

Next, to test the hypothesis that binding modes of dsRBDs are different for conformationally distinct target dsRNAs, TRBP2-dsRBD1 was subjected to NMR-based titration against the four dsRNAs. The primary observation from the ^1^H-^15^N HSQC spectra of TRBP2-dsRBD 1 recorded with the increasing concentration from 0.05 to 0.25 equivalents of RNAs suggested that the interaction of the dsRNAs with the protein led to a broadening of peripheral peaks in the HSQC for the residues in the structured regions of the protein (Figure 3). Such broadening, observed in our study, can be explained by either the presence of an intermediate exchange (at the NMR measurement time scale) between the apo-and the RNA-bound state of the protein or by a decrease in the intrinsic transverse (T_2_) relaxation time of the dsRBD-dsRNA complex. Overall transverse relaxation time (as measured using 1D ^1^H-NMR based T_2_ experiment of free protein and protein-RNA complex) decreased from ~180 ms (in RNA free state) to ~160 ms (in RNA-bound state) suggesting that the change in intrinsic T_2_ may not be the sole cause of line broadening. Therefore, we concluded that the intermediate exchange between free and RNA-bound state of the protein possibly leads to a decrease in apparent T_2_ causing linebroadening in the dsRBD spectra measured at the experimental conditions mentioned above.

**Figure 3:**
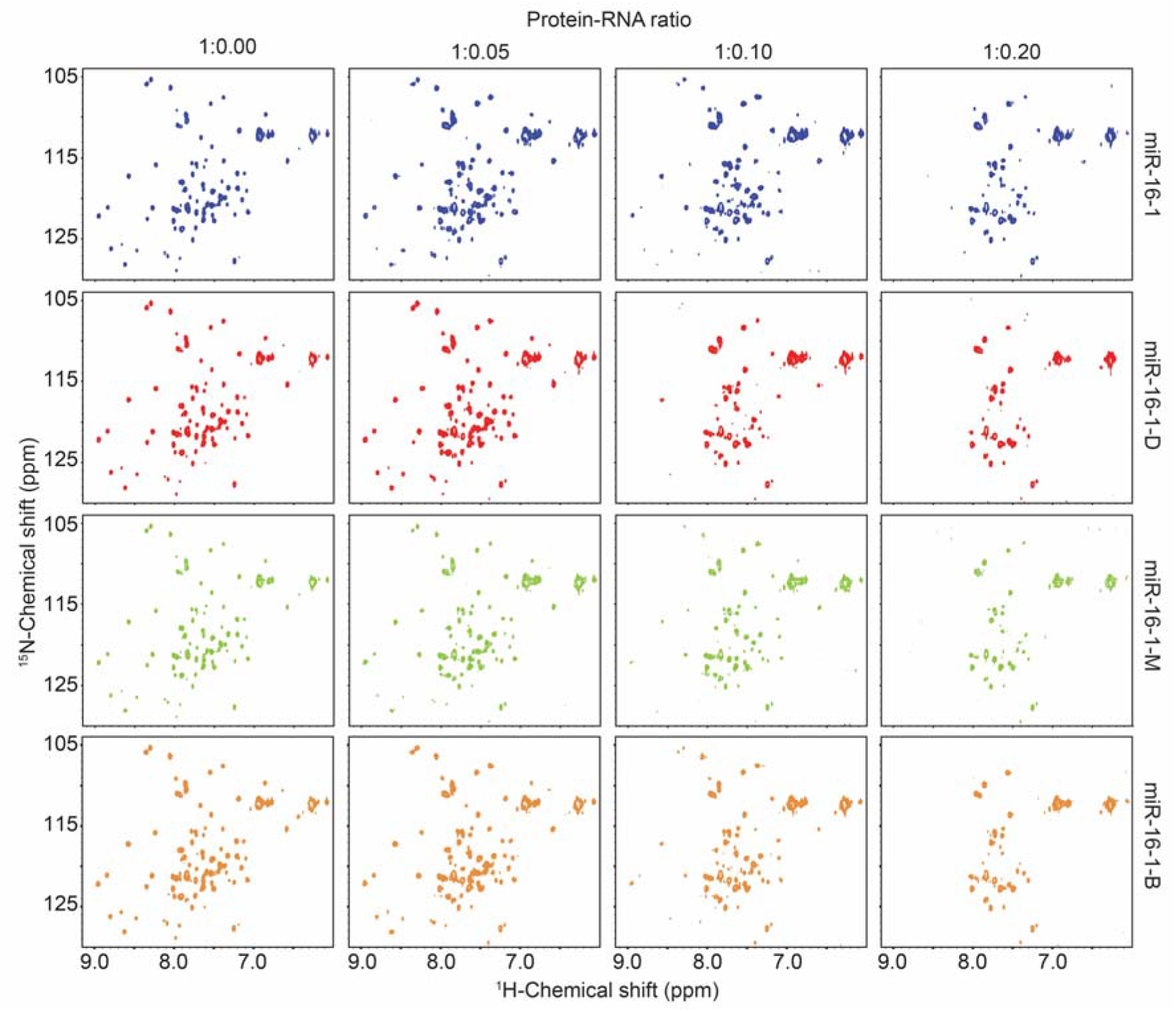
Titration of ^15^N-labeled TRBP2-dsRBD1 against the four dsRNAs as followed by ^1^H-^15^N HSQC experiment. Protein-RNA ratio has been mentioned on the top and is varied from 1:0 to 1:0.2 (Protein:RNA) molar equivalents. Name of the dsRNA has been mentioned on the side. Each successive addition of dsRNA showed a broadening of NMR peaks in the ^1^H-^15^N HSQC spectrum.

As the concentration of the dsRNA was increased to 0.25 equivalents of dsRBD, all the peaks from the residues in the structured regions disappeared with a gradual decrease in the intensity of the peaks (Figure S2). The number and location of the residues showing line-broadening at each successive titration-point (Figure 3 and Figure 4), and the apparent rate of the line-broadening in the four titrations (Figure S2) were found to be different, thereby suggesting that the set of residues of the TRBP2-dsRBD1 undergoing intermediate timescale dynamics is distinct with conformationally distinct dsRNA binding partners. Further, mapping of the residues having line-broadening with increasing RNA concentration showed that the residues involved in the intermediate exchange were present all along the backbone of dsRBD and not localized in the RNA-binding regions (Figure 4) suggesting allosteric role of the broadened residues in the binding event. All the residues broadened in the presence of dsRNAs did exhibit ms-μs timescale dynamics observed in the apo form of the protein (38). No significant change in the intensities was observed for the unstructured terminal residues (M1-L16 except C14, and S99-L105) (Figure 4 and Figure S2). Chemical shift perturbations calculated at 1:0.1 ratio of TRBP2-dsRBD1 to dsRNAs (Figure S3), indicated no significant change in the peak position due to binding as expected for a binding event involving an intermediate exchange between the substrate and the ligand.

**Figure 4:**
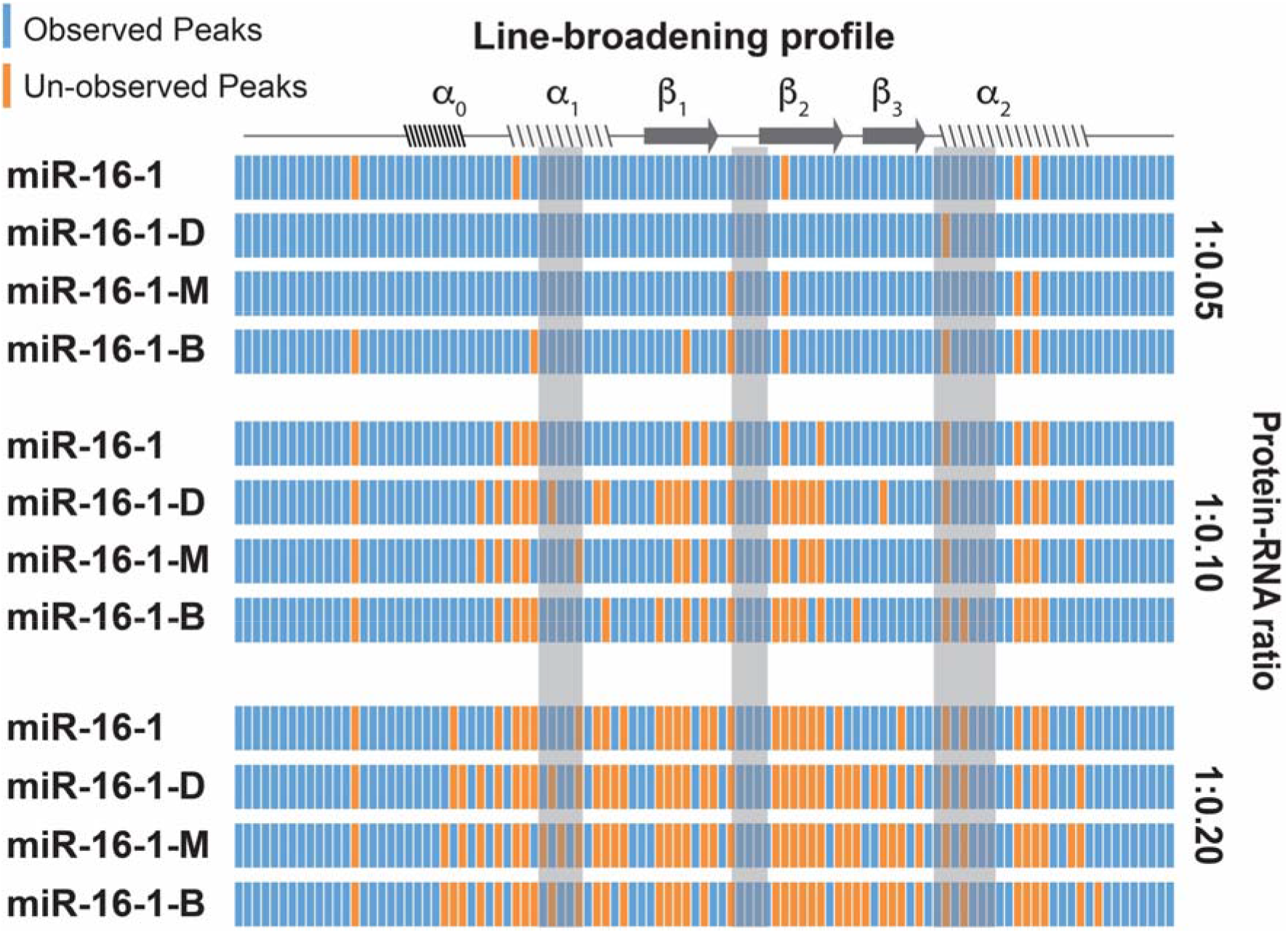
Line-broadening profile obtained from ^1^H-^15^N HSQC of TRBP2-dsRBD1 upon addition of dsRNAs in four different titrations shown in figure 3. Each row corresponds to residues of TRBP2-dsRBD1, where a blue bar represents an observed peak, and an orange bar represents a peak that is disappeared upon addition of addition of dsRNA. Names of the dsRNAs have been mentioned to the left of the corresponding rows. Protein-to-RNA ratio has been mentioned to the right of the rows. Secondary structure of TRBP2-dsRBD1 has been drawn on the top, and the three RNA-binding regions of the protein have been marked with grey vertical bars.

The subtle differences in binding interface of the TRBP2-dsRBD1 as observed in NMR-based titration experiments (Figure 3 and Figure 4) go hand-in-hand with differences perceived in the thermodynamic signatures (ΔH and TΔS) measured by ITC experiments (Figure 2).

### Interaction of dsRBD with a smaller dsRNA

NMR-probed interaction between the dsRBD and the four dsRNAs used in this study suggested an intermediate exchange, which could be ascribed to either between apo- and RNA-bound state of the protein, or to the sliding motion of the protein on the dsRNA as reported earlier (25, 26). To probe this interaction further, we designed a 10-mer dsRNA (D10RNA) sequence derived from miR-16-1-D (Figure 5A). Results of an ITC-based titration (Figure S4) between D10RNA and the two dsRBDs is shown in the figure 5B that illustrates a similar ΔG values of −7.7 and −9.1 kcal/mol for the two dsRBDs but a very different ΔH and TΔS values (Table S2). Interestingly, ^1^H-^15^N HSQC based titration of the TRBP2-dsRBD1 against D10RNA may represent the static binding of dsRBD to dsRNA as has been reported earlier in FRET-based studies for small dsRNA (25). While in the titration of larger dsRNAs, the peak broadening could be observed at very low protein:RNA ratio (1:0.1) (Figure 3), in the titration of shorter dsRNA, the peak broadening started only at 1:1 protein:RNA ratio (Figure S5). A plot of ^1^H-^15^N HSQC peak intensity vs D10-RNA concentration showed that the RNA-induced line-broadening is not localized at RNA-binding regions only and is spread along the backbone (Figure S6), and is not seen at the terminals. Chemical shift perturbations, however, remained of the same magnitude as was seen for the larger RNA sequences (Figure S7).

**Figure 5:**
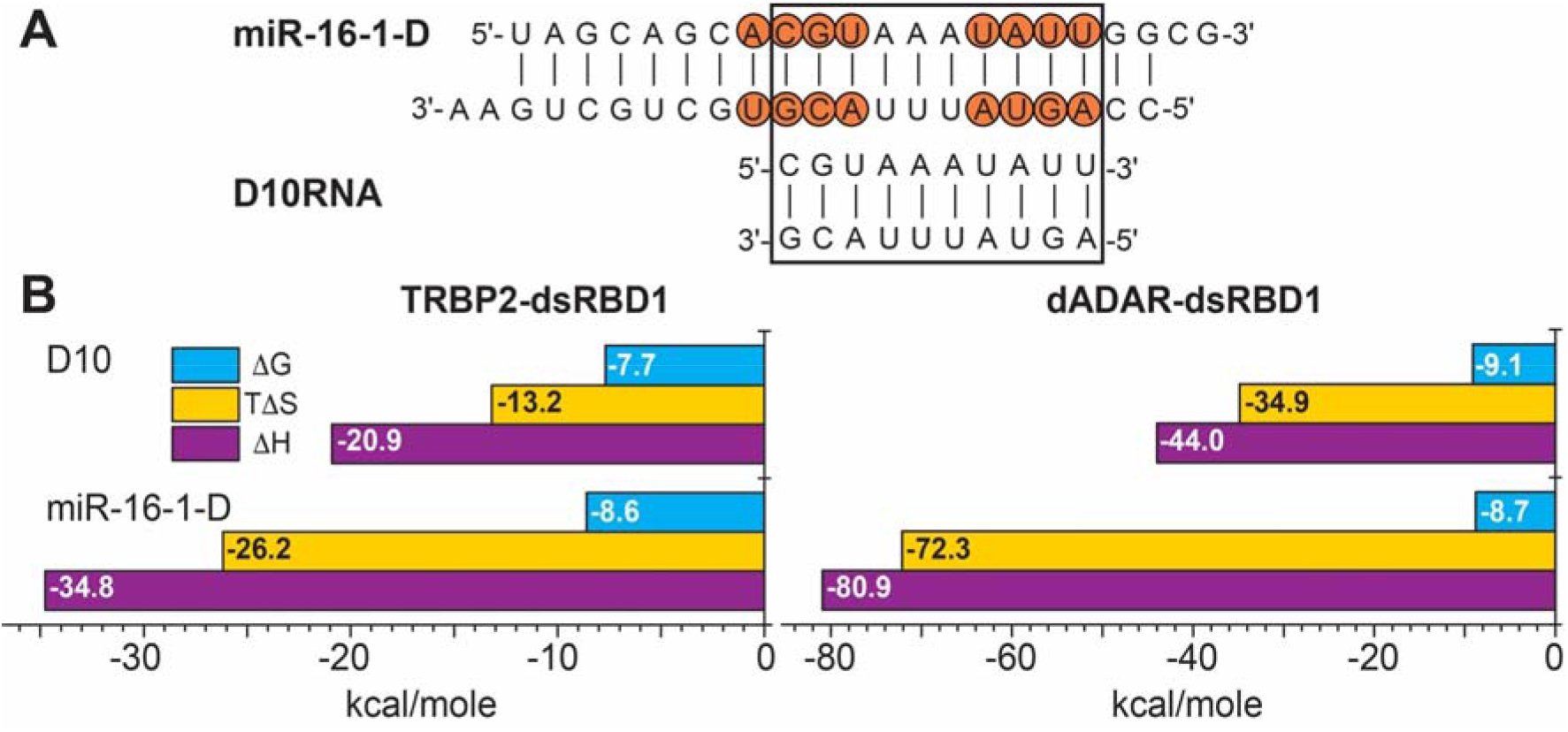
(A) Smaller dsRNA sequence (D10RNA) designed out of miR-16-1-D. Overlapping sequence between the two dsRNAs has been marked with a box. (B) Thermodynamic parameters (ΔG, ΔH, and TΔS) plotted as horizontal bars in the units of kcal/mole as obtained from ITC-based titration of TRBP2-dsRBD1, and of dADAR-dsRBD1 against the D10RNA. Values obtained for titration of TRBP2-dsRBD1 and dADAR-dsRBD1 against miR-16-1-D are also plotted at the bottom for direct comparison with D10RNA.

### Dynamical perturbations in TRBP2-dsRBD1 upon binding with dsRNA

The presence of all the peaks in the D10RNA-bound dsRBD ^1^H-^15^N HSQC spectrum allowed us to probe the dynamics of TRBP2-dsRBD1 in the presence of D10RNA, which could not be done in the presence of larger dsRNAs due to extensive line-broadening. Fast dynamics data (*R*_1_, *R*_2_, [^1^H]-^15^N nOe) could be measured for 58 residues of TRBP2-dsRBD1 bound to D10RNA as against for 69 residues for apo-dsRBD (38). Overall profile for the relaxation parameters (*R*_1_, *R*_2_, [^1^H]-^15^N nOe) was found to be strikingly similar between apo- and dsRNA-bound state (Figure S8) except for a marginal dip in *R*_1_ and *R*_2_ values for the core residues of the protein (highlighted by a horizontal red line in Figure S8) (38). It was concluded from the data that the D10RNA binding did not perturb the overall fast dynamics of the protein.

We then probed the slower microsecond timescale dynamics as was measured for apo-dsRBD using CPMG relaxation dispersion (CPMG-RD) (Figure S9) and heteronuclear adiabatic relaxation dispersion (HARD) NMR experiments (Figure S10) and compared the same with the ones measured for apo-dsRBD (38). CPMG-RD plots showed that while most of the residues displayed no relaxation dispersion (as was also seen for the apo-TRBP2-dsRBD1 (38), relaxation dispersion was seen for the following residues, namely, Y48, L50, L51, A53, A82, A88, E89, and L95 (Figure S9). Though the dispersion in relaxation rates was observed, due to large errors in the *R_2eff_* values, dynamics parameters could not be extracted.

As for apo-form of the protein, HARD NMR experiments were performed to observe dynamics in dsRBD in presence of the RNA substrate. The plot of *R*_1ρ_ and *R*_2ρ_ data for apo- and D10RNA-bound TRBP2-dsRBD1 plotted against residue numbers showed dispersion in the relaxation rates, suggesting presence of dynamics at ms-μs timescale in RNA-bound state similar to apo-state of the dsRBD (38). The HARD NMR relaxation data was further analyzed using a two-state model as was done for the apo-dsRBD (38, 51) and exchange parameters (*k*_ex_, pB, and Δω) were extracted and compared with those for the apo-dsRBD. Interestingly, in the RNA-binding regions, the exchange appeared to have quenched upon RNA-binding (disappearance of red balls and appearance of blue/green balls in figure 6). The residues which were exchanging with *k*_ex_ > 50000 s^−1^ in the apo- dsRBD, are now exchanging with lower *k*_ex_ values. Intriguingly, a few residues which were exchanging with *k*_ex_ < 50000 s^−1^ are now found to undergo exchange at an increased *k*_ex_ as shown by an appearance of red balls at new locations in the RNA-bound protein (Figure 6). The perturbation in *k*_ex_ upon binding to D10RNA was closely monitored by obtaining the difference in *k*_ex_ values between bound and apo-dsRBD (Figure 7A). A positive Δ*k*_ex_ would suggest an increase in *k*_ex_ upon binding to D10RNA, while a negative value would suggest a quench of the exchange process. We have color coded the Δ*k*_ex_ > 50000 s^−1^ on the protein backbone to highlight the residues which underwent a drastic change upon D10RNA binding (Figure 7B). It was fascinating to observe that the residues where the exchange process seemed to quench upon D10RNA binding (marked with cyan in Figure 7B and 7C) are in close spatial proximity to the ones where *k*_ex_ has increased in D10RNA-bound state (marked with red in Figure 7B and 7C). For example, in region 1, exchange in Q36 (belonged to dsRNA-binding region 1) was suppressed, while that in L34 and G39 exchange was induced upon binding with D10RNA. Similarly, region 2 (not an RNA-binding region, a *k*_ex_ > 50000 s^−1^ observed in this region was attributed to allosteric role of residues to provide the stability to the protein in apostate) was highlighted with a quench of exchange in T67 and V68, while a simultaneous increase of exchange in V47 and T71. In region 3, residues lying in or near to RNA-binding regions (N61 and A82) exhibited a quench, while a neighboring residue F62 showed an increase in *k*_ex_.

**Figure 6:**
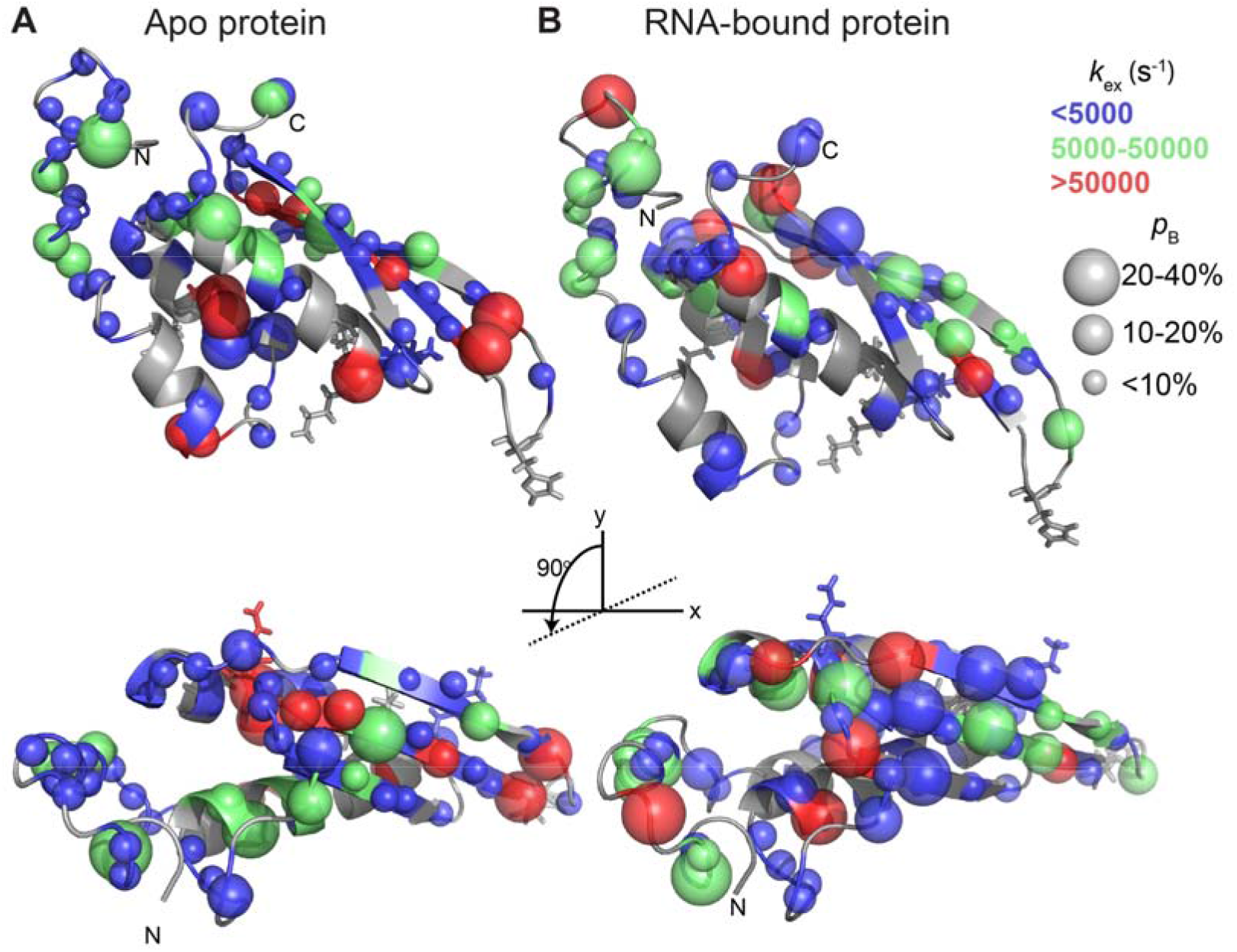
Conformational exchange along the backbone of the two dsRBDs. Mapping of dynamics parameters, rate of exchange between ground state and excited state (*k_ex_*), and population of the excited state (*p_B_*) as obtained by the ‘geometric approximation method’ from HARD experiment, on the tertiary structure of (A) apo-TRBP2-dsRBD1 and (B) D10RNA-bound TRBP2-dsRBD1. The color code has been used to highlight the distribution of *k_ex_* values whereas the size of the sphere highlights the variation in the *p_B_* values across protein backbone. The RNA binding residues have been shown in stick mode.

**Figure 7:**
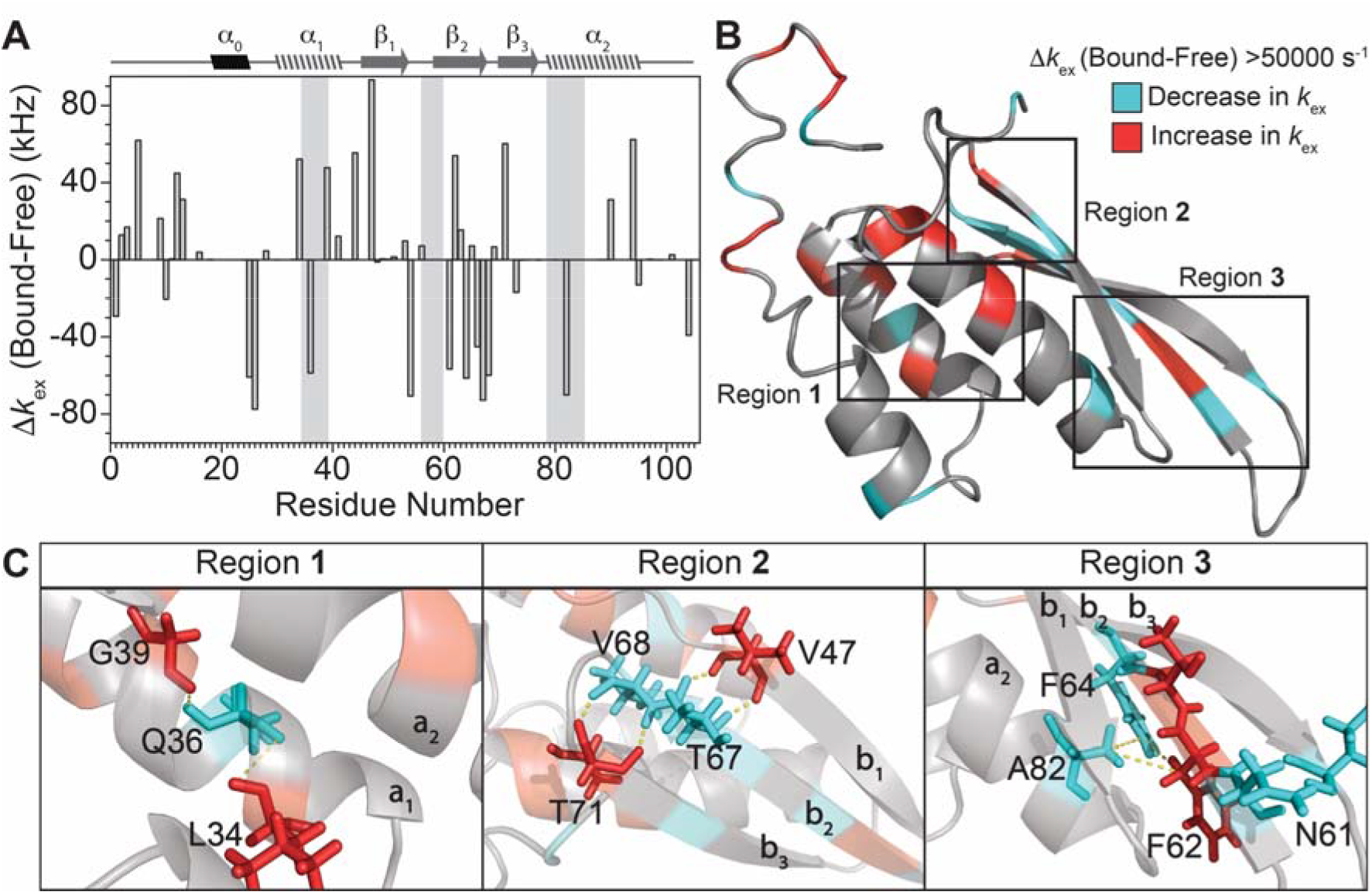
Summary of perturbations in conformational exchange in TRBP2-dsRBD1 upon binding with D10RNA. (A) Plot of Δ*k*_ex_ (D10RNA-bound – Apo-TRBP2-dsRBD1, as obtained from HARD NMR experiments) against residue numbers. (B) An increase in *k*_ex_ upon D10RNA binding is marked in red and a decrease in *k*_ex_ has been marked in cyan on the backbone of TRBP2-dsRBD1. (C) Three regions of the backbone are zoomed in to highlight the spatial proximity between residues undergoing increase in *k*_ex_ with those undergoing decrease in *k*_ex_ upon D10RNA binding.

## Discussion

In an attempt to understand the role of protein dynamics being played in the conformational adaptation required for dsRBDs to target and bind dsRNAs of varying shapes, two dsRBDs from two different proteins from two species – TRBP2-dsRBD1 from *Homo sapiens* and dADAR-dsRBD1 from *Drosophila melanogaster* – were employed against four different dsRNAs with varying shapes. The primary sequence of the four dsRNAs differed in single nucleotide deletion or mutation, that resulted into overall variation in global conformation (Figure 1). ITC-based titration of the four dsRNAs with the two dsRBDs showed very intriguing results (Figure 2 and Figure S1). While the ΔG of binding reaction remained similar (~8 kcal/mol) for all four dsRNAs with the two dsRBDs, as expected due to only subtle changes in dsRNA sequences, a great deal of change was observed in ΔH and TΔS values of the binding reaction from one dsRNA sequence to another, and from one dsRBD to the other. For each binding partner, a distinct combination of ΔH and TΔS values were obtained from ITC data analysis. Since the net change in free energy of the binding were observed to be similar in all cases, the system depicts a well-studied case of Enthalpy/Entropy compensation (55–57, 63, 64). The results also show that the favorable enthalpic change during binding is compensated by the loss of entropy in going from apo-protein to the protein-RNA complex and the process is dominated by the enthalpic term. A negative value of TΔS in ITC-based titration could be ascribed to the unfavorable entropic cost of formation of a dynamically constrained dsRBD-dsRNA complex. The enthalpic contribution can originate from the multiple point of contacts that well-conserved RNA-binding residues of dsRBD make with dsRNA through hydrogen bonding, van der Waals interactions, etc. Though the dsRBDs interact with dsRNAs via known binding sites, differences in ΔH observed for conformationally different dsRNAs highlights the variations in binding modes. The differences in ΔH is also manifested in the number of protein molecules binding to a particular dsRNA as the binding sites obtained from ITC measurements. We may thus conclude that the binding mode of a given dsRBD to a given dsRNA is dependent on the overall shape of the dsRNA.

Furthermore, the titration of TRBP2-dsRBD1 against the four dsRNAs probed by ^1^H-^15^N HSQC experiment corroborated the finding that the binding mode of dsRBD to a given dsRNA is unique. No conclusions about structural changes in dsRBD upon dsRNA-binding could be drawn from the chemical shift perturbations as 1) the overall perturbations were very minimal (Figure S3), and 2) extensive line-broadening was observed in the presence of dsRNAs (even at 1:0.05 protein:RNA ratio, Figure 3). However, a gradual line-broadening upon addition of dsRNAs exhibited a unique pattern of residues (Figure 4 and Figure S2) that were being broadened at each successive titration point for a given dsRNA further suggesting that each dsRNA was interacting with the dsRBD with a unique binding mode. The line-broadening observed was also suggestive of an intermediate exchange (53, 65–67) that could be attributed to either between apo- and dsRNA-bound dsRBD, or to the sliding motion of the dsRBD on a dsRNA as have been reported earlier (25, 26). To test the presence of sliding motions, a shorter dsRNA (derived from one of the four dsRNAs used above but containing only 10 base pairs), D10RNA, was employed (Figure 5A). An ITC-based titration of D10RNA with the two dsRBDs again showed similar ΔG values with a unique combination of ΔH and TΔS values (Figure 5B and Figure S4). The number of protein binding sites with D10RNA were found to be 2.5±0.1 and 2.4±0.1 for TRBP2-dsRBD1 and dADAR-dsRBD1, respectively. A ^1^H-^15^N HSQC titration for TRBP2-dsRBD1 against D10RNA did not show the line-broadening (up until 1:1 protein:RNA ratio, Figure S5 and Figure S6) as was seen for larger dsRNA suggesting static dsRBD binding with D10RNA.

Fast (ps-ns timescale) dynamics was measured on TRBP2-dsRBD1 complexed with D10RNA and was compared with that measured earlier on apo-TRBP2-dsRBD1(38). No significant change in the relaxation parameters were observed other than customary marginal decrease in *R*_1_ and *R*_2_ parameters due to increase in overall size. The comparison of slow (microsecond timescale) dynamics between apo-and D10RNA-bound TRBP2-dsRBD1, however, was quite exciting. On one hand, quenching of microsecond timescale dynamics was observed upon RNA-binding in the RNA-binding residues; conformational exchange seemed to get induced at other locations. Similar switch in the dynamics was also observed in the residues that were proposed to have allosteric role in RNA-binding. Further, the analysis of the positions of these residues in the structure show that the residues undergoing decrease and increase in *k*_ex_ are in close spatial proximity, thereby suggesting a relay of conformational exchange from one site to the other site upon dsRNA-binding. Yamashita et al have shown that the interaction of dsRBD of TRBP with dsRNA does not lead to significant change in the structure of dsRBD (41), while a report by Acevedo et al. have shown that the interaction with a dsRBD also does not affect the structure of the RNA (68). Distribution of the dynamics in the apoprotein and its perturbation upon binding with a dsRNA suggested that the conformational exchange observed in the apo-protein helps it to interact with dsRNA. The apparent relay of conformational exchange from one site to the other site upon dsRNA-binding thus suggests that the protein is highly adaptive in nature. The shift of the positions of dynamic residues from the apo-protein to the RNA-bound protein suggest the importance of intrinsic dynamics to adapt to target a variety of dsRNA-shapes.

## Conclusions

In conclusion, the investigations from the current study show that dsRBDs recognize dsRNAs in a shape-dependent manner and the binding modes of such recognition are unique for topologically distinct dsRNAs. Further, the interaction between dsRBD-dsRNA studied here exhibits a well-studied case of enthalpy-entropy comensation, where the overall process is enthalpy-driven. A quench of intrinsic conformational exchange, which was observed in apo-dsRBD, was seen along with the induction of conformational exchange at new sites in close spatial proximity to quench sites. This apparent relay of conformational exchange from one site to another suggested the adaptive nature of the dsRBD, a critical feature required to target topologically distinct dsRNAs.

## Author Contribution

J.C. and H.P. conceived the study, designed dsRNA sequences, approach to dynamically characterize the model dsRBDs in RNA-bound state, and wrote the paper. H.P. performed all the experiments and analysed the data in active discussions with J.C.

## Acknowledgements

Authors thank Prof. Jennifer Doudna (University of California, Berkeley) for the TRBP plasmids and Prof. Frederic H.T. Allain (ETH Zurich) for ADAR-plasmid. Authors also acknowledge High Field NMR facility at IISER-Pune (co-funded by DST-FIST and IISER Pune) and the High-Field NMR facility at IIT Bombay. Authors are also grateful to Dr. Sinjan Choudhary and Mr. Mayuresh for providing access to ITC facility at UM-DAE CEBS, Mumbai. This work was supported by funding from Indian Institute of Science, Education and Research, Pune; Department of Biotechnology, Govt. of India [No. BT/PR24185/BRB/10/1605/2017]; and extramural funding from the Science and Engineering Research Board (SERB), Govt. of India [EMR/2015/001966]. J.C. thanks the International Collaborative Research Grant obtained from Institute of Protein Research, Osaka University for the machine time on 950 MHz. Authors also acknowledge Prof. Gianluigi Veglia (University of Minnesota, Minnesota) and Dr. Fa-An Chao (National Cancer Institute, Maryland) for active discussions while setting up HARD experiments and data analysis. Authors thank Dr. Arnab Mukherjee and Dr. T.S. Mahesh at IISER Pune for helpful discussion as Research Advisory Committee members for HP. Authors also thank Dr. Shilpy Sharma (Savitribai Phule Pune University, Pune) for active discussions during the data analysis and writing of the manuscript. HP is thankful to IISER Pune for the fellowship. HP is thankful to CSIR (India) [No. TG/10468/19-HRD] and Infosys Foundation [No. IISER-P/InfyFnd/Trv/139] for Travel support to present part of this work at Gordon Research Conference on Computational Aspects of Biomolecular Research, Switzerland, June 9-14, 2019.

## Supplementary Information

Supplementary Information (SI) available.

