## Supplementary Material for "Role of protein dynamics in enthalpy-driven recognition of topologically distinct dsRNAs by dsRBDs"

Harshad Paithankar^1^ and Jeetender Chugh^1,2,^*

^1^Department of Chemistry, and ^2^Department of Biology, Indian Institute of Science Education and Research (IISER), Dr. Homi Bhabha Road, Pashan, Pune 411008, India

**Running Title**: Protein-dynamics and dsRBD-dsRNA interaction

**Figure S1**

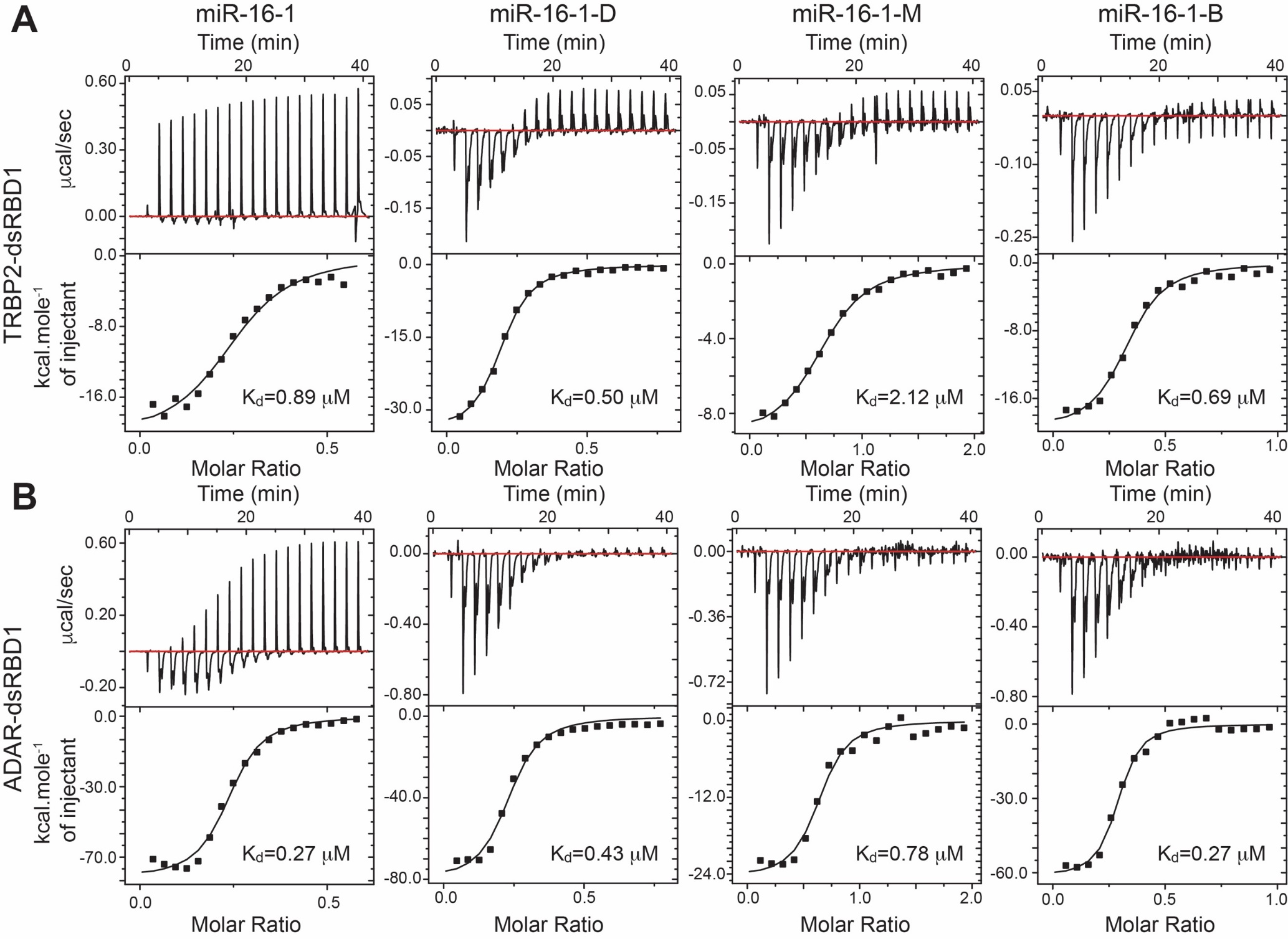

Figure S1: Isothermal Titration Calorimetry binding profiles of (A) TRBP2-dsRBD1, and (B) dADAR-dsRBD1 against four dsRNAs mentioned in Figure 1. The name of the corresponding dsRNA has been mentioned on the top. Within each panel, the top part shows raw data of heat change per injection as a function of time and the bottom part shows processed data of heat change as a function of molar ratio of RNA:Protein. K_d_ values for each titration obtained from data fitting (see methods section) have been mentioned in the inset.

**Figure S2**

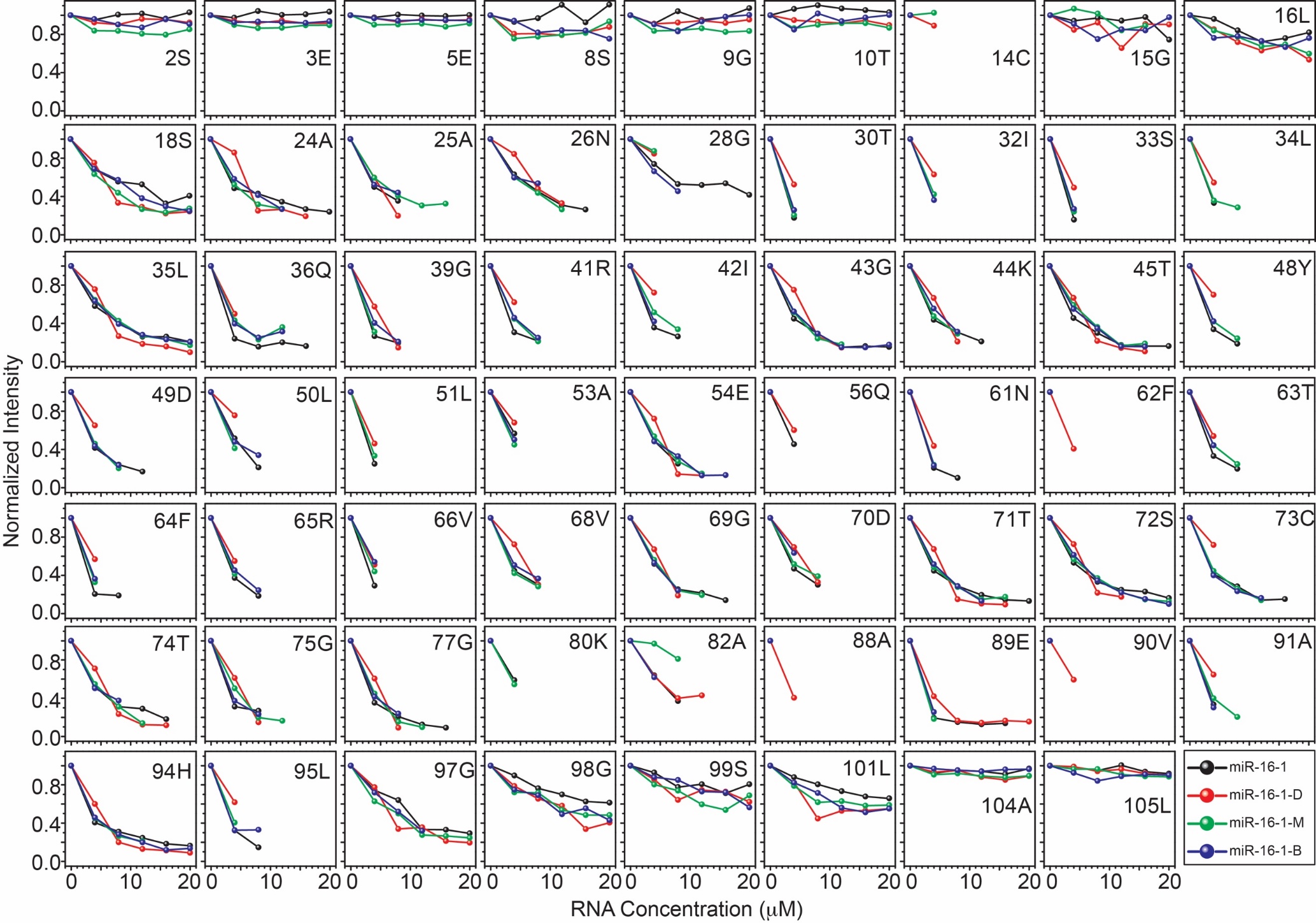

Figure S2: NMR peak intensity from ^1^H-^15^N HSQC spectra of TRBP2-dsRBD1, depicted in Figure 3, have been plotted against RNA concentrations. The color code used to mark distinct dsRNAs has been mentioned in the bottom right panel.

**Figure S3**

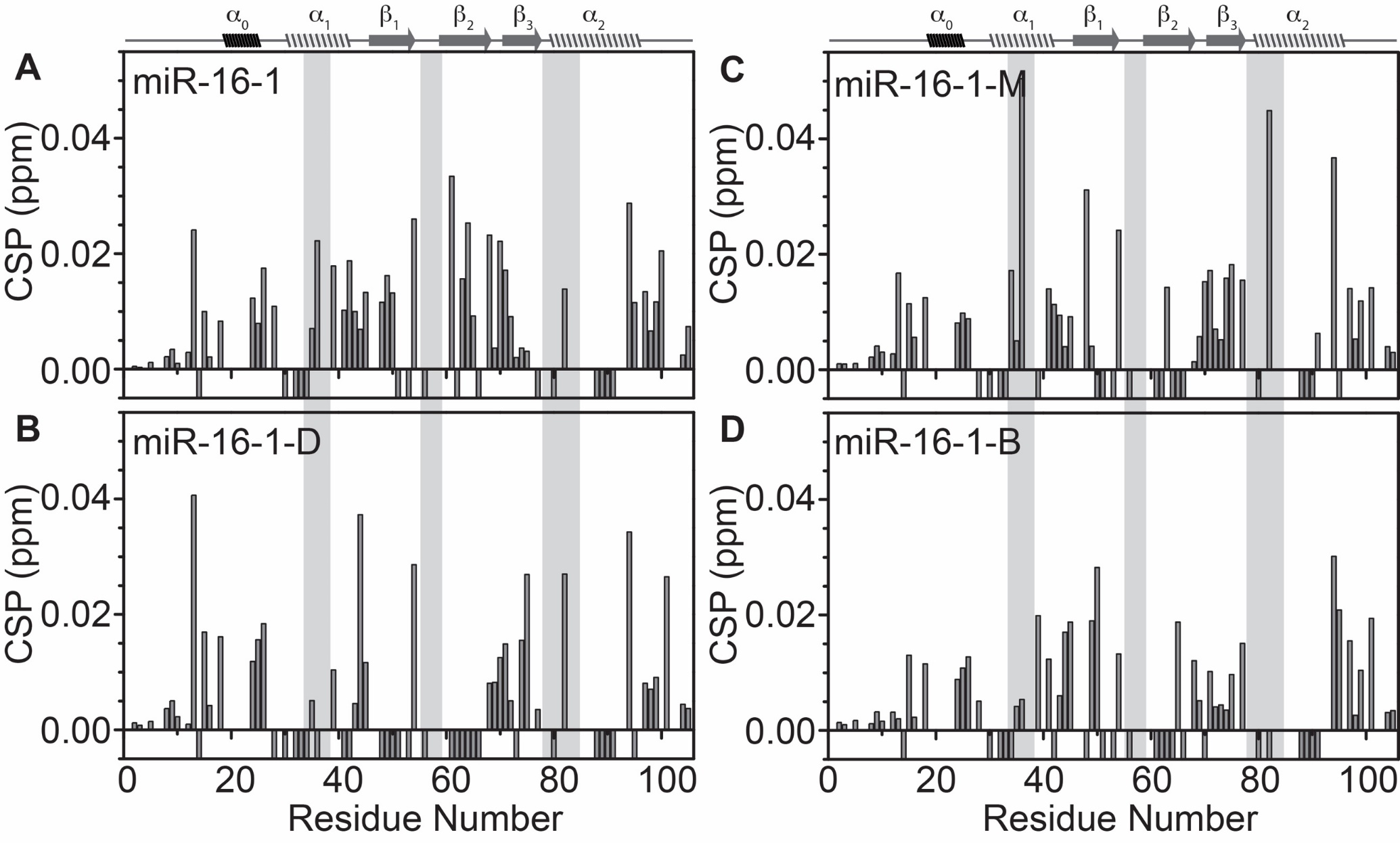

Figure S3: Chemical shift perturbations (CSP in ppm) in ^1^H and ^15^N chemical shift dimensions from ^1^H-^15^N HSQC spectra of TRBP2-dsRBD1 have been plotted against residue number for (A) miR-16-1, (B) miR-16-1-D, (C) miR-16-1-M, and (D) miR-16-1-B at Protein:RNA concentration ratio of 1:0.1. The secondary structure of TRBP2-dsRBD1 has been mentioned on the top, and the RNA-binding region of the protein has been marked in grey vertical columns. Negative CSP values indicate broadened peak for which CSP could not be measured.

**Figure S4**

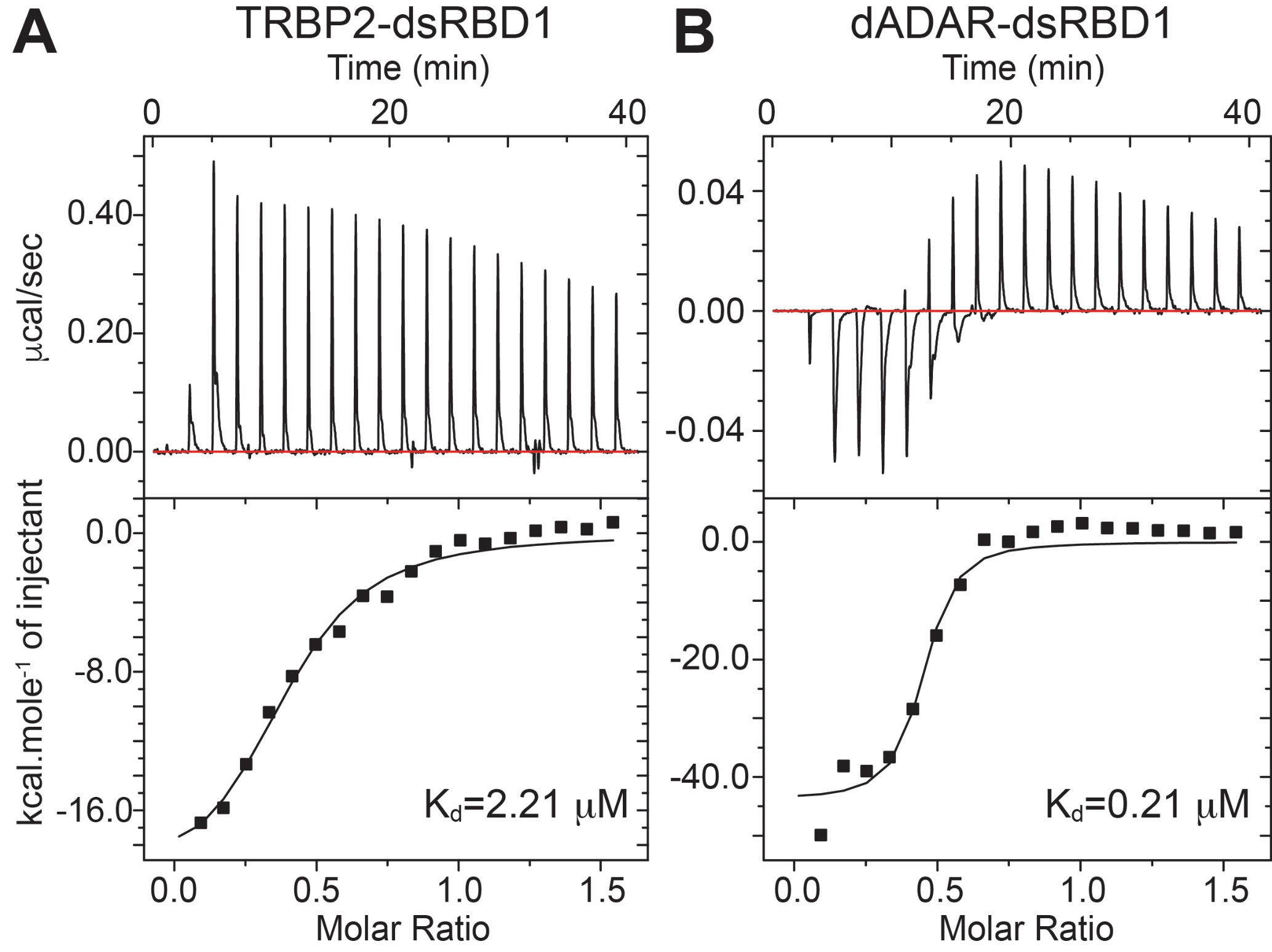

Figure S4: Isothermal Titration Calorimetry binding profiles of (A) TRBP2-dsRBD1, and (B) dADAR-dsRBD1 against the smaller dsRNA construct, D10RNA, as mentioned in figure 5A. Within each panel, the top part shows raw data of heat change per injection as a function of time and the bottom part shows processed data of heat change as a function of molar ratio of RNA:Protein. K_d_ values for each titration obtained from data fitting (see methods section) have been mentioned in the inset.

**Figure S5**

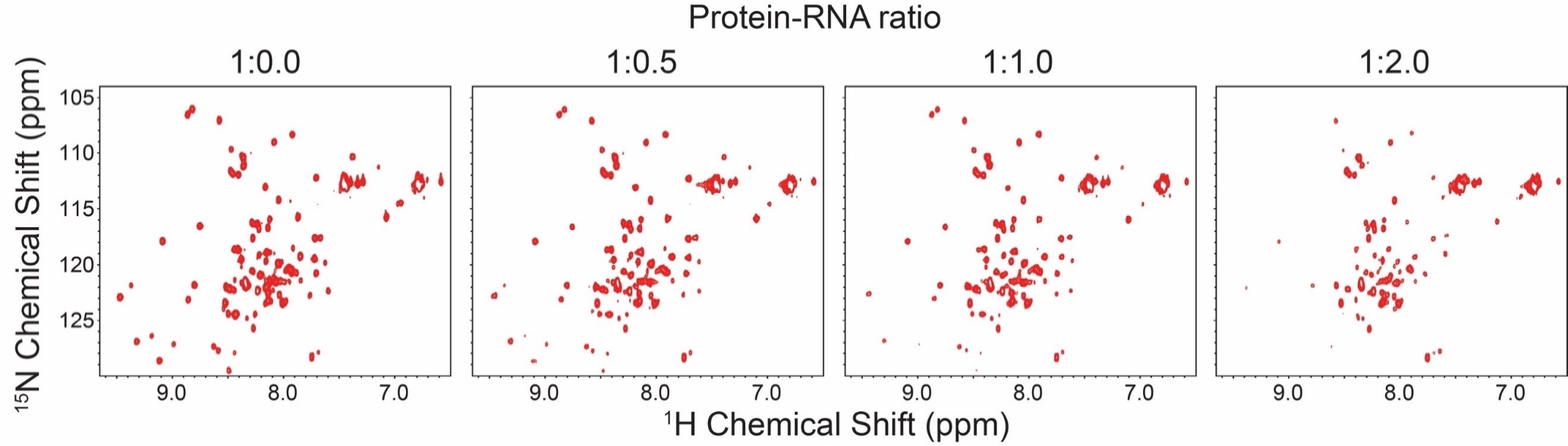

Figure S5: Titration of ^15^N-labeled TRBP2-dsRBD1 against the D10RNA as followed by ^1^H-^15^N HSQC experiment. The protein-RNA ratio has been mentioned on the top and is varied from 1:0 to 1:2 (Protein:RNA) molar equivalents. Unlike in Figure 3, line-broadening only started at 1:1 for this dsRNA.

**Figure S6**

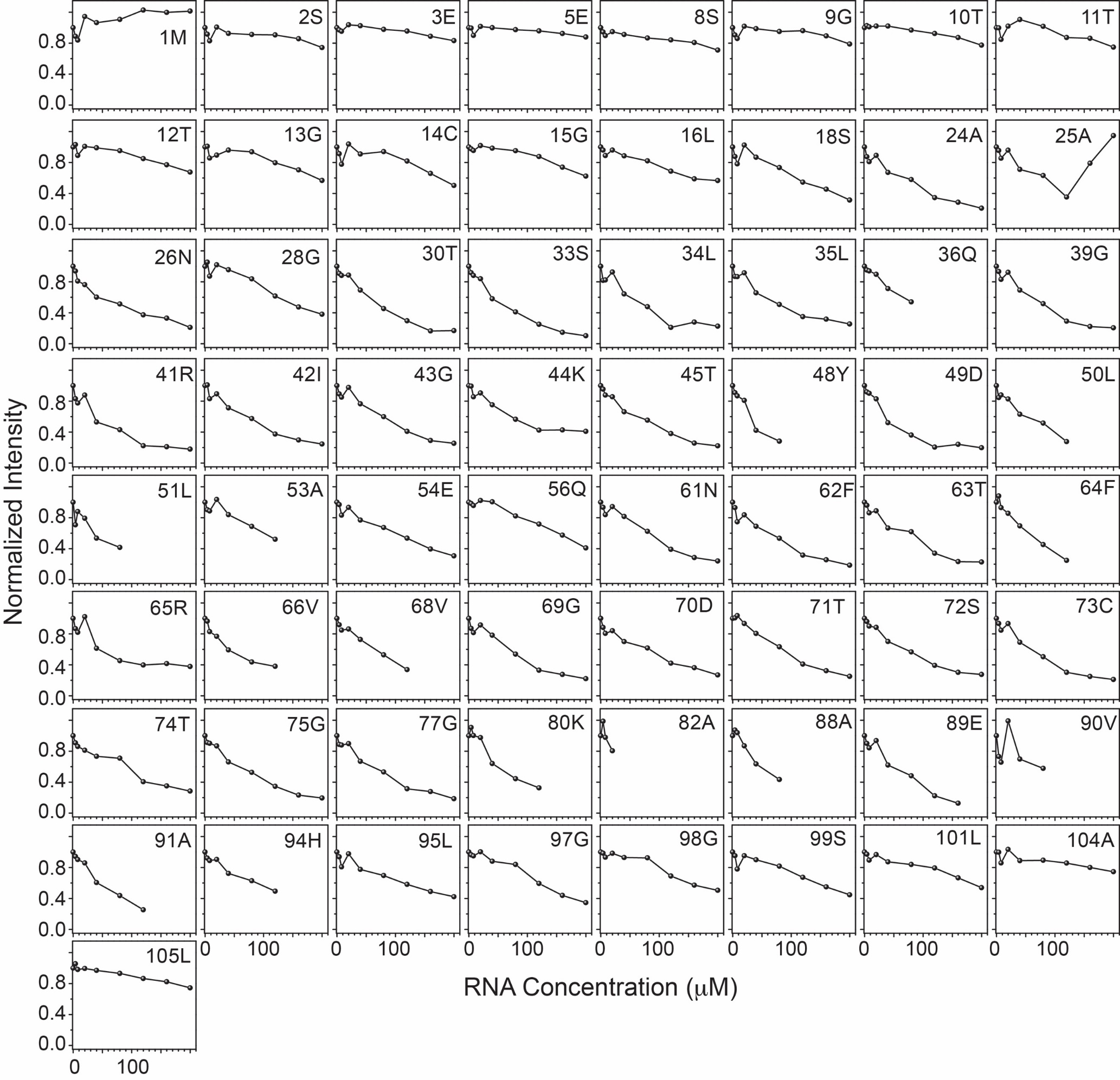

Figure S6: NMR peak intensity from ^1^H-^15^N HSQC spectra of TRBP2-dsRBD1, depicted in figure S5, has been plotted against RNA concentrations.

**Figure S7**

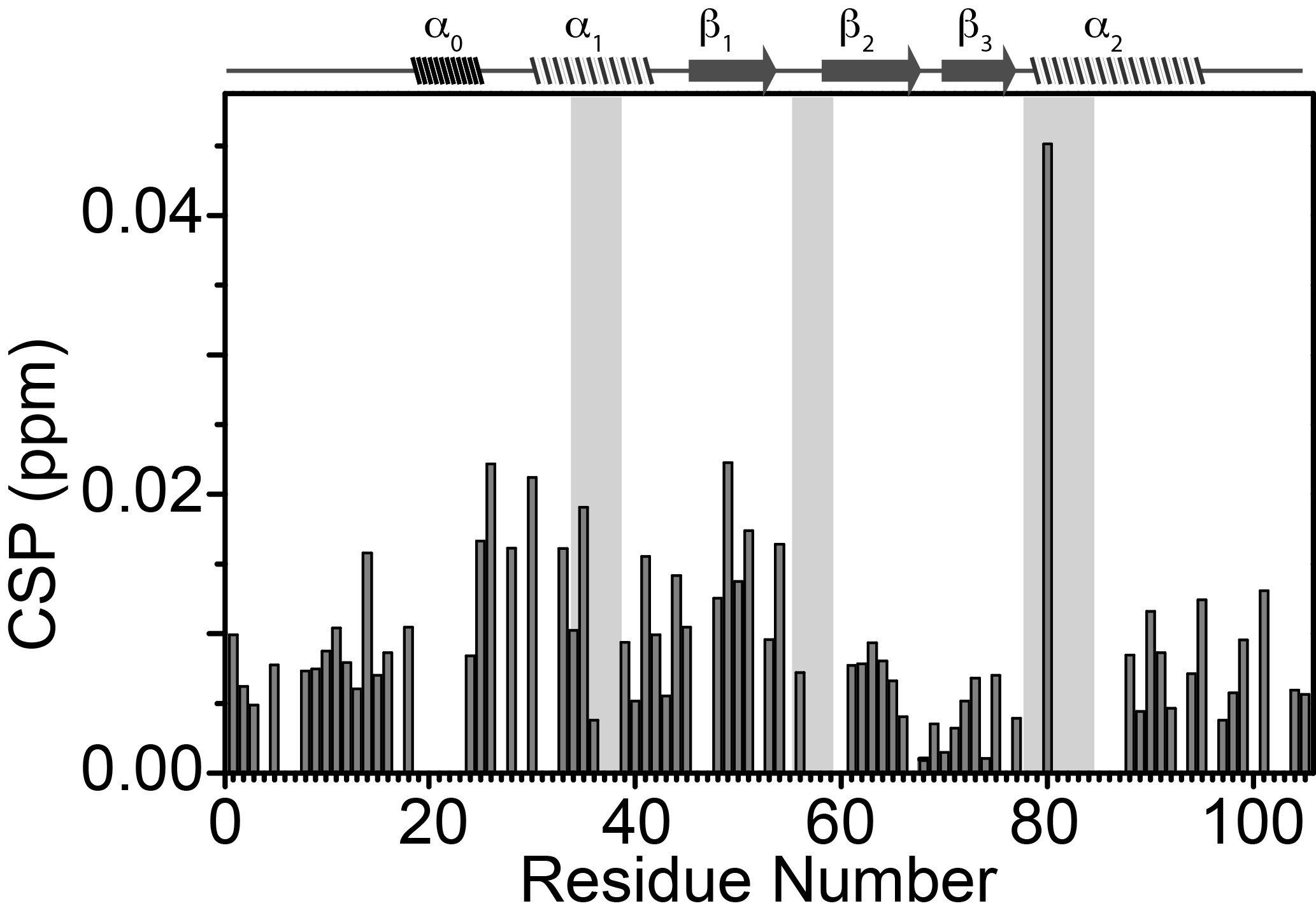

Figure S7: Chemical shift perturbations (CSP in ppm) in ^1^H and ^15^N chemical shift dimensions from ^1^H-^15^N HSQC spectra of TRBP2-dsRBD1 plotted against residue number for D10RNA at Protein:RNA concentration ratio of 1:1. The secondary structure of TRBP2-dsRBD1 has been mentioned on the top, and the RNA-binding region of the protein has been marked in grey vertical columns.

**Figure S8**

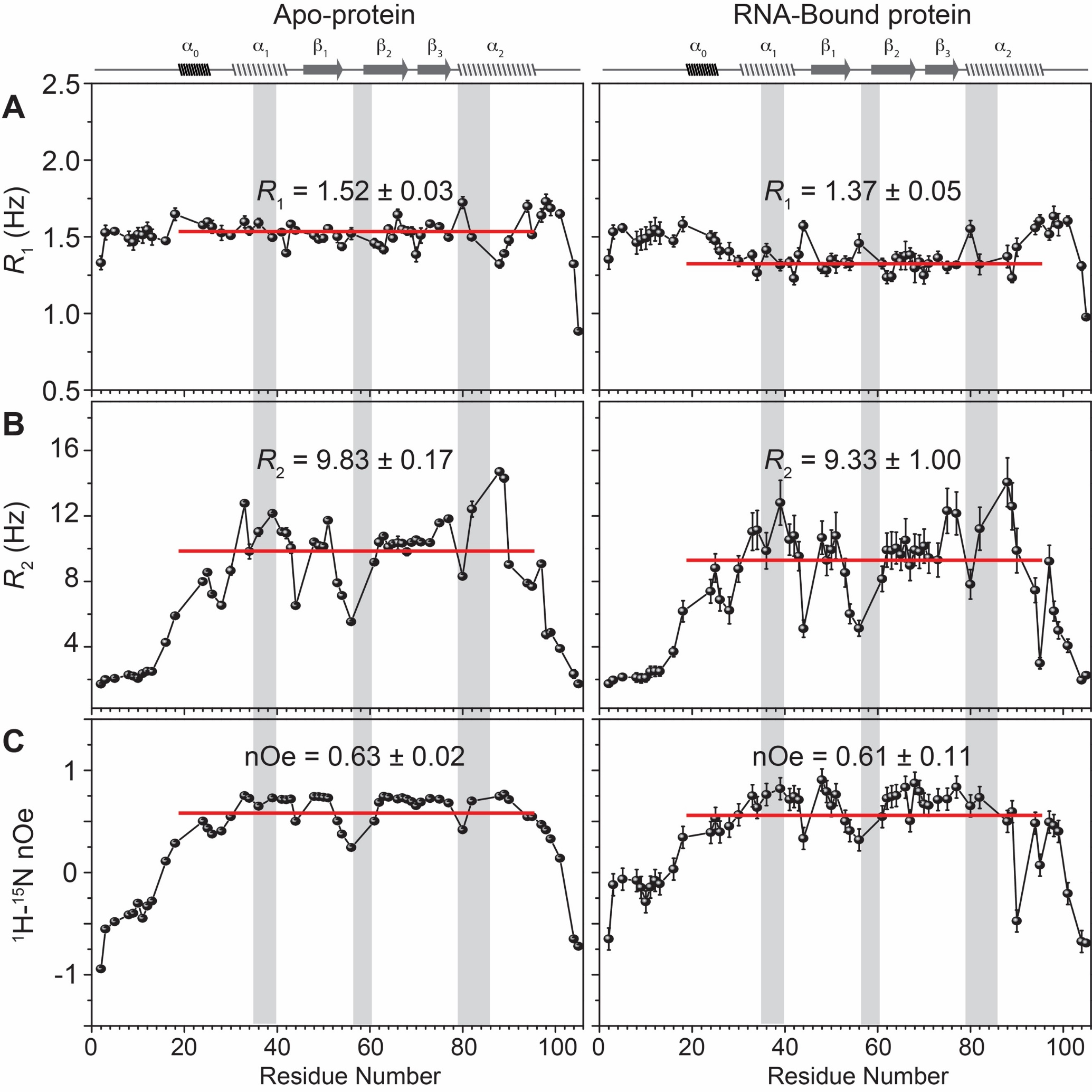

Figure S8: Plot of relaxation parameters (A) *R*_1_, (B) *R*_2_, and (C) [^1^H]-^15^N nOe against residue numbers for apo-protein (left panel, reported earlier in our study (1)), and for D10RNA-bound TRBP2-dsRBD1 (right panel). The secondary structure of TRBP2-dsRBD1 has been mentioned on the top, and the RNA-binding region of the protein has been marked in grey vertical columns.

**Figure S9**

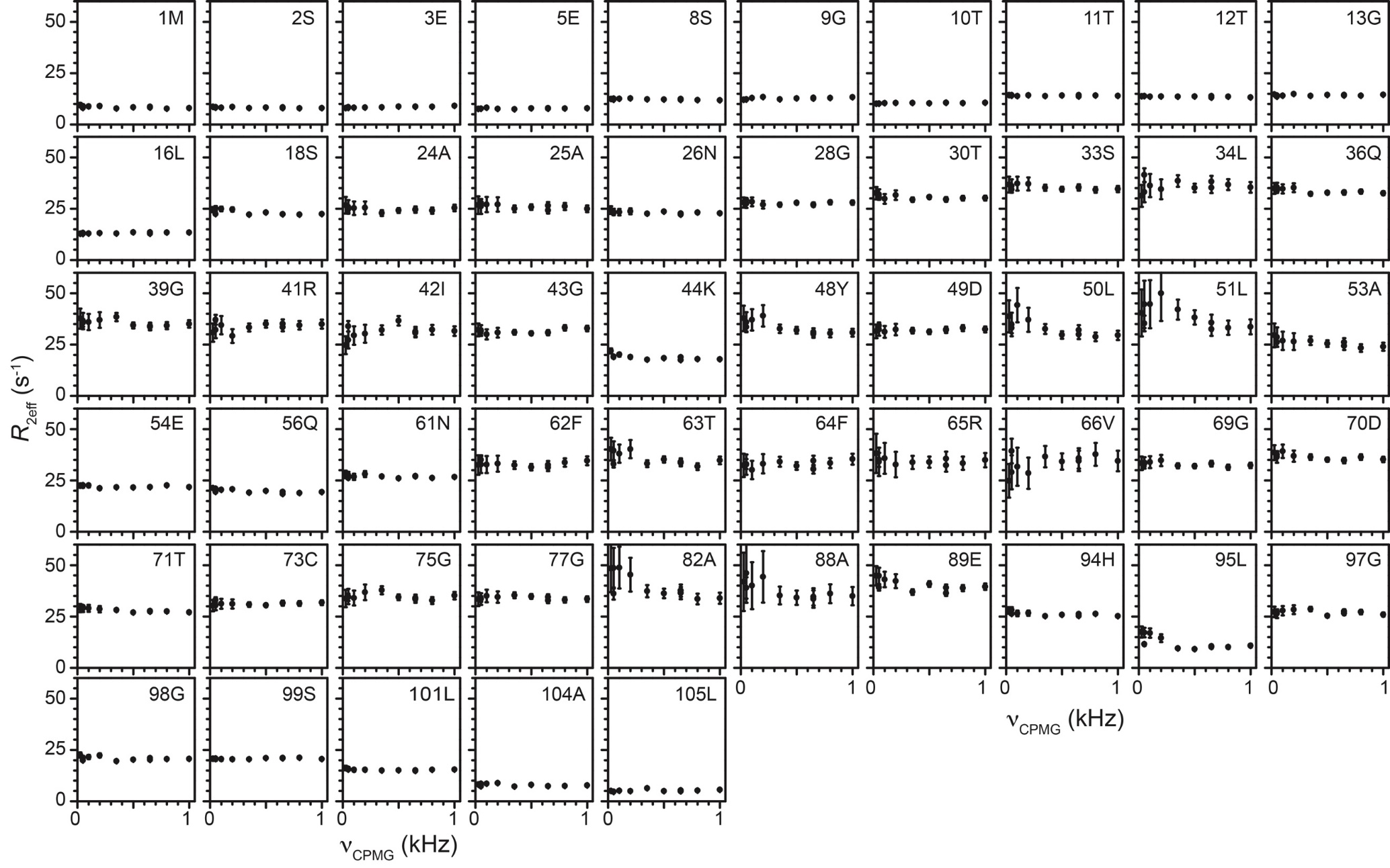

Figure S9: *R*_2eff_ rates as obtained from CPMG relaxation dispersion experiments plotted against the CPMG frequency for all the non-overlapping peaks of D10RNA-bound TRBP2-dsRBD1.

**Figure S10**

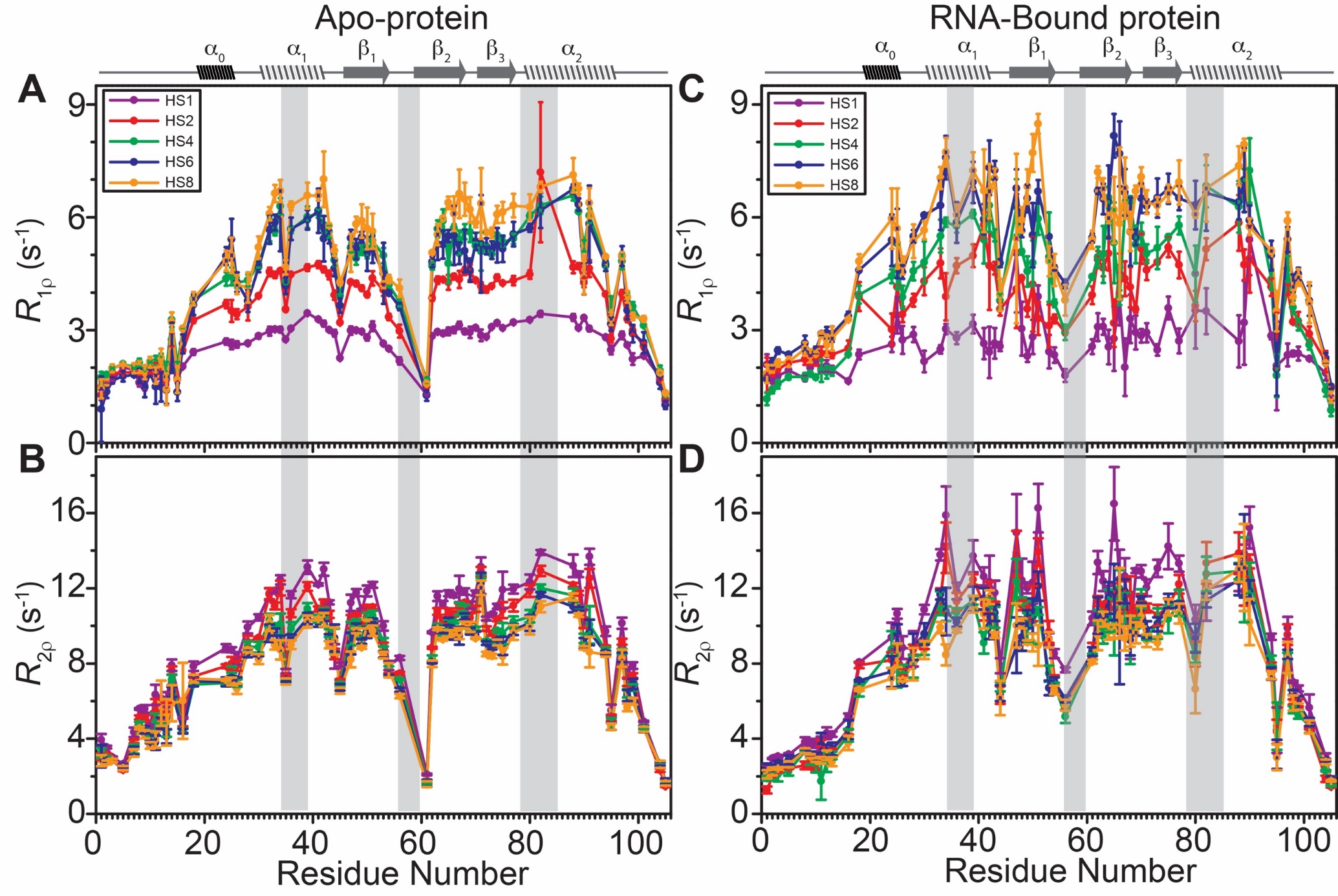

Figure S10: Dispersion in relaxation rates for TRBP2-dsRBD1. (A) *R*_1ρ_ and (B) *R*_2ρ_ relaxation rates derived from the adiabatic relaxation dispersion experiments plotted against residue number as a function of adiabatic pulse stretching factors (color coded for n=1,2,4,6, and 8) for apo-TRBP2-dsRBD1 as reported earlier in (1). (C) *R*_1ρ_ and (D) *R*_2ρ_ relaxation rates from the adiabatic relaxation dispersion experiments plotted for D10RNA-bound TRBP2-dsRBD1.

**Table S1**

Table S1: RNA sequences used to study interaction with dsRBDs

| **Name** | **Strand** | **Sequence (5ʹ 🡪 3ʹ)** |
| --- | --- | --- |
| miR-16-1 | Guide | 5ʹ- UAGCAGCACGUAAAUAUUGGCG -3ʹ |
|  | Passenger | 5ʹ- CCAGUAUUAACUGUGCUGCUGAA -3ʹ |
| miR-16-1-D | Guide | 5ʹ- UAGCAGCACGUAAAUAUUGGCG -3ʹ |
|  | Passenger | 5ʹ- CCAGUAUUUACGUGCUGCUGAA -3ʹ |
| miR-16-1-M | Guide | 5ʹ- UAGCAGCACGUAAAUAUUGGCG -3ʹ |
|  | Passenger | 5ʹ- CCAGUAUUAACGUGCUGCUGAA -3ʹ |
| miR-16-1-B | Guide | 5ʹ- UAGCAGCACGUAAAUAUUGGCG -3ʹ |
|  | Passenger | 5ʹ- CCAGUAUUAACGUGCUGCUGAA -3ʹ |
| D10RNA | Guide | 5ʹ- UUAUAAAUGC -3ʹ |
|  | Passenger | 5ʹ- GCAUUUAUGA -3ʹ |

**Table S2**

Table S2: Thermodynamic parameters calculated for dsRBD:dsRNA interaction calculated from fitting of Isothermal Titration Calorimetry data to one-set of sites binding model

| **RNA sequences** | **TRBP2-dsRBD1** | **dADAR-dsRBD1** |
| --- | --- | --- |
| ***K_d_* (μM)** | | |
| miR-16-1 | 0.89 ± 0.21 | 0.27 ± 0.05 |
| miR-16-1-D | 0.50 ± 0.04 | 0.43 ± 0.10 |
| miR-16-1-M | 2.12 ± 0.22 | 0.78 ± 0.24 |
| miR-16-1-B | 0.69 ± 0.13 | 0.27 ± 0.07 |
| D10RNA | 2.21 ± 0.51 | 0.21 ± 0.10 |
| **Number of Binding sites** | | |
| miR-16-1 | 3.9 ± 0.1 | 4.3 ± 0.1 |
| miR-16-1-D | 5.2 ± 0.1 | 4.4 ± 0.1 |
| miR-16-1-M | 1.5 ± 0.0 | 1.6 ± 0.1 |
| miR-16-1-B | 3.0 ± 0.1 | 3.6 ± 0.1 |
| D10RNA | 2.5 ± 0.1 | 2.4 ± 0.1 |
| ***ΔH* (kcal.mole^-1^)** | | |
| miR-16-1 | -20.6 ± 1.1 | -69.0 ± 1.8 |
| miR-16-1-D | -34.8 ± 0.6 | -80.9 ± 3.2 |
| miR-16-1-M | -9.4 ± 0.2 | -24.6 ± 1.1 |
| miR-16-1-B | -19.8 ± 0.7 | -61.8 ± 2.0 |
| D10RNA | -20.9±1.6 | -44.0 ± 2.2 |
| ***TΔS* (kcal.mole^-1^)** | | |
| miR-16-1 | -12.4 ± 1.0 | -59.9 ± 1.8 |
| miR-16-1-D | -26.2 ± 0.6 | -72.1 ± 3.1 |
| miR-16-1-M | -1.6 ± 0.2 | -16.3 ± 0.9 |
| miR-16-1-B | -11.4 ± 0.6 | -52.8 ± 1.9 |
| D10RNA | -13.2 ± 1.4 | -34.9 ± 2.0 |
